# Genetic background diversifies mutational paths but converges on membrane depolarization for bleomycin resistance in yeast

**DOI:** 10.64898/2026.07.28.741370

**Authors:** Yan Shuai, Guihong Lai, Ludong Yang, Haiqing Zeng, Li Zhang, Jia-Xing Yue, Jing Li

**Author notes:** Corresponding authors: Li Zhang, Jia-Xing Yue, Jing Li. Yan Shuai and Guihong Lai contributed equally to this work.

## Abstract

Drug resistance evolution is shaped by both selective agent and genetic background in which resistant mutations arise, but most experimental evolution studies rely on a single genetic background. Here we performed parallel experimental evolution across six genetically diverged *Saccharomyces cerevisiae* strains, propagating 48 replicate populations for ∼225 generations under the selection of DNA-damaging chemotherapeutic bleomycin. Over 70% of the evolved populations acquired resistance with the magnitude and repeatability of adaptation varying markedly across genetic backgrounds. Whole-genome sequencing showed that bleomycin elevated single-nucleotide and insertion-deletion mutation rates, and induced background-specific aneuploidies and structural variants, including Ty-element-mediated translocations and tandem duplications. Notably, mutation rates varied up to 10-fold among backgrounds, with the most commonly-used lab strain exhibiting the highest mutagenic susceptibility, underscoring that evolutionary insights derived from single-background models may not universally generalize across a species. Despite this background-dependent genomic heterogeneity, we identified recurrent loss-of-function mutations in *PMA1* and *PTK2* that arose independently across nearly all genetic backgrounds and conferred strong resistance. Functional and biophysical assays revealed that these mutations impair Pma1p H^+^-ATPase activity and depolarize the plasma membrane, thereby limiting the cellular uptake of the positively charged drug. Our findings demonstrate that while the mutational routes to drug resistance are heavily shaped by genetic background, a convergent, background-independent mechanism centered on membrane potential regulation can dominate phenotypic adaptation. This work underscores the necessity of integrating genetic diversity into experimental evolution paradigm and nominates plasma membrane potential as an important lever for modulating chemotherapeutic efficacy.

## Introduction

Darwinian evolution drives phenotypic adaptation not only in natural ecosystems but also in the biomedical contexts, most consequentially in the emergence of drug resistance. Despite substantial therapeutic advances, resistance remains one of the greatest obstacles in modern medicine. Understanding the genomic and phenotypic dynamics by which individuals adapt is therefore essential both for dissecting the mechanisms underlying resistance and for developing strategies to overcome it.

Experimental evolution of microorganisms offers a powerful approach for studying the evolutionary dynamics in real time [1,2]. Replicate populations, from a common ancestor under controlled condition for many generations, allow phenotypes to be tracked longitudinally while samples are archived at defined timepoints for in-depth analysis. This approach has enabled genotype-phenotype links to be drawn for resistance mechanisms with direct relevance to mammalian and human cells [3–5].

Experimental evolution of budding yeast (*Saccharomyces cerevisiae*), typically using standard lab strains like the BY background, have provided fundamental insights into the dynamics and mechanisms of adaptation [6–8]. Despite these advances, relying on such single-background experimental setups frequently obscure the profound influence of genetic background. Our previous studies demonstrated that the mutation rates, initial responses to stresses and the subsequent evolutionary trajectories towards adaptation all vary among strains with distinct backgrounds [9–11]. Consistent with these findings, recent studies using bacteria and yeast also underscore the pivotal role of genetic background in shaping both phenotypic and genomic evolutionary dynamics [12–15]. The pre-existing genetic variants dictate the initial stress response, and their interaction with the type and duration of selection shapes the acquisition of *de novo* mutations, together determining the ultimate phenotypic outcome [16,17]. More importantly, these dynamics extend far beyond the lab settings. Patients with distinct genetic backgrounds often show markedly different responses to the same treatment, varying substantially in both the kinetics and the evolutionary routes through which resistance emerges [18,19]. Therefore, incorporating multiple genetic backgrounds into the study design of experimental evolution is essential for understanding how drug resistance arises and evolves, as well as dissecting the underlying mechanisms.

Bleomycin is a radiomimetic glycopeptide antibiotic. It has been used to treat cancer for decades and remains a standard therapy for several cancer types like testicular cancer. Bleomycin induces cytotoxicity through formation of a DNA-bound metal complex (e.g. Fe^2+^) that catalyzes the generation of reactive oxygen species (ROS), causing single-strand and double-strand DNA breaks. Prior high-throughput genetic screen and short-term experimental evolution in yeast lab strains (BY background) have identified resistance mechanisms involving disrupted drug transport and loss of vacuolar detoxification [20–22]. However, whether these mechanisms are conserved across genetically diverse individuals or represent idiosyncrasies of the lab strain in which they were discovered remains unknown. The extent to which bleomycin-induced mutagenesis itself varies with genetic background is also largely unknown.

To address these questions, we performed parallel experimental evolution of six genetically diverged *S. cerevisiae* strains under bleomycin selection, propagating 48 replicate populations for ∼225 generations. By tracking the phenotypic dynamics, we found that over 70% of the evolved lineages acquired resistance to bleomycin, though the magnitude and consistency of adaptation varied considerably by genetic background. Whole-genome sequencing of ancestral and evolved populations allowed us to estimate mutation rates and comprehensively profile genomic alterations, both among replicate lineages and across genetic backgrounds. This revealed substantial background-dependent variation in mutagenesis alongside recurrent structural rearrangements unique to specific backgrounds. Despite this heterogeneity, we identified *de novo* mutations in *PMA1* and *PTK2* that arose repeatedly across nearly all genetic backgrounds. Using targeted genetic and functional assays, we show that these mutations converge on a shared mechanism of resistance, disrupting plasma membrane potential in a manner that limits cellular uptake of bleomycin. Together, our results demonstrate how genetic background shapes the mutational landscape of drug resistance evolution while still permitting a conserved, background-independent route to adaptation.

## Results

### Multi-background parallel evolution of *S. cerevisiae* under bleomycin stress

To investigate how individuals with distinct genetic backgrounds adapt to stress, we used six homozygous diploid *S. cerevisiae* strains, including lab strain BY4743 (BY), Wine/European strain DBVPG6765 (WE), West African strain DBVPG6044 (WA), North American strain YPS128 (NA), Sake strain Y12 (SA), and Malaysian strain UWOPS03-461.4 (MA), for parallel adaptive evolution under bleomycin treatment (**Fig. 1A**). The five natural strains (WE, WA, NA, SA, and MA) harbor 51,664 to 91,029 SNPs relative to the lab strain (BY), reflecting their deep evolutionary divergence [23]. Prior to initiating the experimental evolution, we profiled the baseline growth of each ancestral strain to determine an optimal bleomycin concentration that is capable of exerting substantial selective pressure without causing complete lethality. We chose a concentration of 12.5 μg/mL, which extended the generation time of the ancestral strains by 11.6% to 95.6% across the six strains (mean = 45%). This pronounced baseline variation underscores substantial phenotypic differences in drug susceptibility of these genetic backgrounds prior to any adaptation (**Fig. S1**).

**Figure 1.**
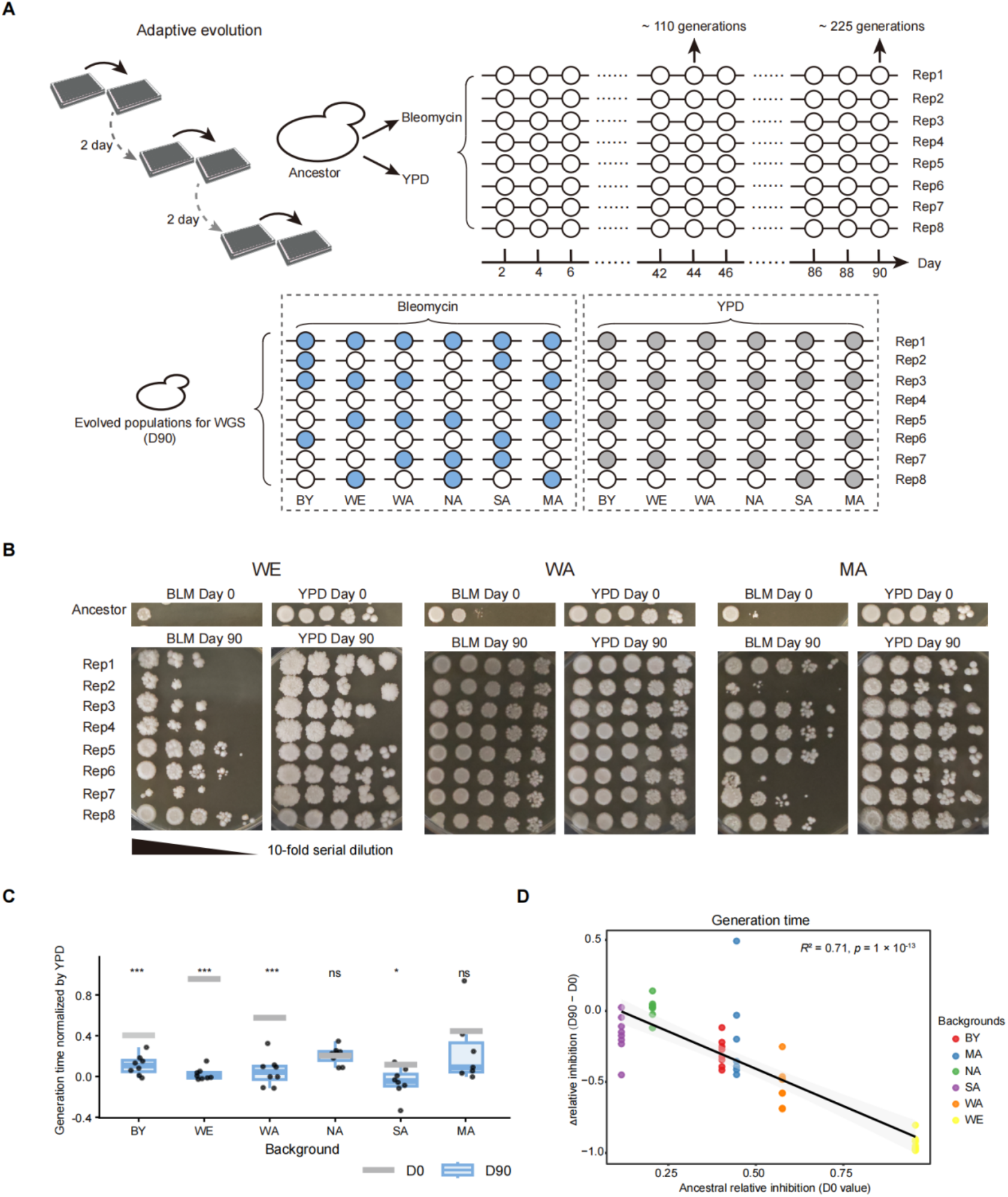
Experimental evolution of bleomycin resistance. **(A)** Six diploid yeast strains with diverse genetic backgrounds independently evolved for 90 days in either YPD alone or YPD supplemented with 12.5 μg/mL bleomycin, with eight replicate populations per background per condition. At day 90, four replicates per condition were sequenced (indicated by filled circles). Populations were designated using the format: “day_strain_condition_replicate” (e.g., “D90_WE_BLM_1” represents replicate 1 of Wine/European strain evolved in bleomycin for 90 days). **(B)** Phenotypic evolution under bleomycin stress. Spotting assays were performed using bleomycin-evolved samples at indicated time points on YPD plates with or without 12.5 μg/mL bleomycin. Ten-fold serial dilutions were spotted from left to right, with eight replicates arranged from top to bottom. Representative results are shown; the complete dataset is available in Fig. S1A. **(C)** Enhanced bleomycin resistance in evolved strains. We normalized the generation time in the presence of drug by that in its absence, calculated as (with BLM − without BLM) / without BLM. Compared to D0, the evolved BY, WE, and WA and SA populations showed a significant reduction in generation time (*** p < 0.001, * p < 0.05, t-test) while NA and MA exhibited no significant difference (ns, p > 0.05). **(D)** Correlation between initial growth inhibition and growth improvement measured by generation time. Growth improvement was calculated as the change in growth inhibition between D90 and D0 strains. Each point represents an independently evolved lineage. Pearson correlation coefficients (R²) and corresponding *P* values are shown in the panel.

Using this optimized concentration, we propagated eight replicate populations per ancestral strain in either bleomycin-supplemented or in drug-free YPD medium for 90 days, corresponding to approximately 225 generations (**Fig. 1A**). The dynamics of growth phenotypes under bleomycin stress were monitored by spotting assay across six timepoints (days 0, 2, 8, 20, 50, and 90). Resistant clones were detectable as early as day 2, suggesting that early-arising variants may have seeded subsequent adaptation of the population. By day 90, more than 70% of the evolved populations exhibited enhanced bleomycin resistance (**Fig. 1B and Fig. S1A**). Our quantitative growth assay further corroborated this observation, revealing a significantly increased fitness under bleomycin treatment for the evolved populations relative to their respective ancestors (**Fig. 1C,** *P* < 0.05, t-test; **Fig. S1B-S1C, Table S1**). The magnitude of fitness improvement was inversely correlated with ancestral fitness (**Fig. 1D, Fig. S1D-S1E**), consistent with diminishing-returns epistasis whereby beneficial mutations confer proportionally smaller gains in already-fitter genetic backgrounds [24].

Despite this general trend toward increased resistance, we observed pronounced variation among genetic backgrounds in both the magnitude and reproducibility of adaptation. For example, while all eight WA replicates evolved near-complete resistance, the MA populations exhibited more heterogeneity with six MA populations achieving modest-to-high resistance whereas the other two remaining substantially sensitive (**Fig. 1B**). Together, these results establish that bleomycin resistance arises predictably under selection, but that ancestral genetic background strongly shapes both the degree and the repeatability of the phenotypic outcome.

### Aneuploidy and structural variation

To characterize bleomycin-induced mutagenesis across genetic backgrounds, we sequenced the ancestors, 48 evolved populations (24 population in bleomycin-supplemented and 24 in drug-free YPD) at day 90, and 27 clones isolated from the bleomycin-evolved populations (**Table S2**). In drug-free YPD, one NA and one SA population exhibited multiple chromosome copy-number alterations, corresponding to a baseline aneuploidy rate of 7.41 × 10^⁻4^ events per line per generation (**Fig. S2A**). Aneuploidy remained infrequent under bleomycin selection, occurring at a rate of 1.30 × 10⁻³ events per line per generation (**Fig. 2A, Fig. S2A)**. Notably, bleomycin-induced aneuploidy was enriched in specific genetic backgrounds, particularly MA and SA. Half of the sequenced MA populations (2/4) carried aneuploidies with multiple chromosome gains, recurrently affecting chromosomes XIII and V. We further validated these findings by sequencing isolated clones (**Fig. S2B**). Similarly, two independently evolved SA populations gained chromosomes VIII and XII. In contrast, we did not observe such frequent aneuploidy occurrences in other genetic backgrounds. These patterns point to background-specific chromosome instability under bleomycin stress. While whether these aneuploidies directly confer resistance remains to be fully elucidated, their independent recurrence strongly hints at a likely adaptive advantage.

**Figure 2.**
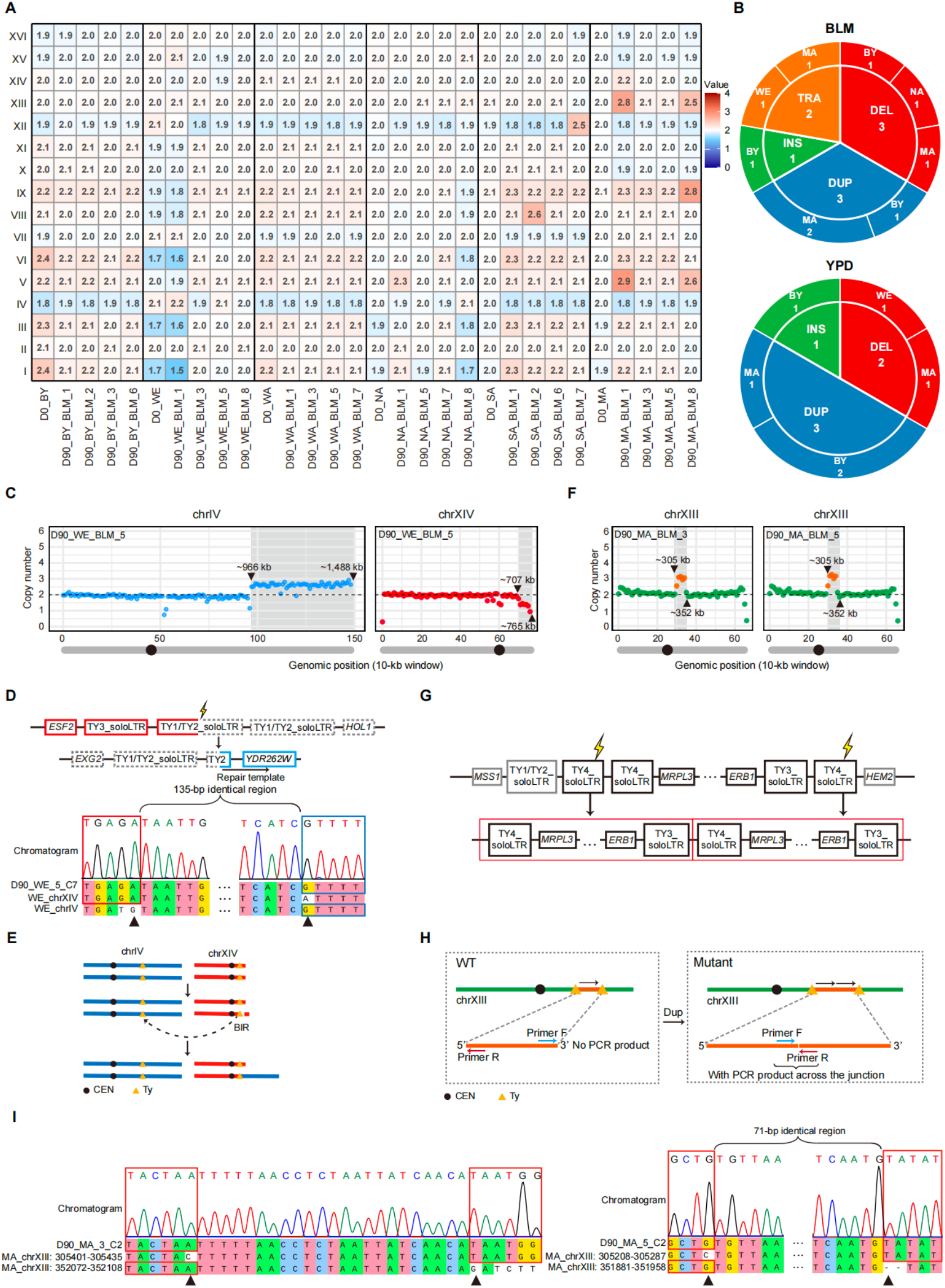
Aneuploidy and structural variation (SV) in bleomycin-evolved populations. **(A)** Chromosome copy number variation (CNV) across evolved populations. Chromosome copy number = 2 × (median depth of the chromosome / median depth of the genome). **(B)** SV number identified in YPD and bleomycin conditions. **(C)** Copy number profiles in 10-kb windows along chrIV and chrXIV for population D90_WE_BLM_5. The shaded regions indicate a segmental terminal gain and loss, respectively. **(D)** Sanger sequencing chromatogram and gene map around the breakpoint in the clone C7 from D90_WE_BLM_5. **(E)** Schematic of the translocation between chrIV and chrXIV. **(F)** Copy number profiles in 10-kb windows along chrXIII for population (left panel) D90_MA_BLM_3 and (right panel) D90_MA_BLM_5, with the shaded regions indicating segmental duplications. **(G)** Gene map surrounding the duplicated region on chrXIII. **(H)** Schematic and primer design for the chrXIII duplication event. **(I)** Sanger sequencing verification of the breakpoints in (left panel) clone C2 from D90_MA_BLM_3 and (right panel) clone C2 from D90_MA_BLM_5. Distinct breakpoint sequences between the two clones from different populations are highlighted.

Beyond whole-chromosome aneuploidies, we next examined whether bleomycin also induced structural variation (SV). While the frequencies of large insertions, deletions and duplications remained low and comparable between bleomycin and the YPD control conditions (**Fig. S3**), we identified two translocation events unique to the bleomycin condition (**Fig. 2B**). Specifically, in population D90_BLM_WE_5, we detected a ∼550 kb terminal gain on the right arm of chromosome IV (chrIV-R), accompanied by a ∼60 kb terminal loss on the right arm of chromosome XIV (chrXIV-R) (**Fig. 2C**), suggesting an unbalanced translocation where the chrIV terminal segment was appended to the right end of chrXIV. This rearrangement was supported by multiple chimeric paired-end sequencing reads spanning the translocation junction. Notably, the chrXIV breakpoint region contains several retrotransposon fragments (a Ty3 solo-LTR and Ty1/Ty2 solo-LTR), while the corresponding locus on chrIV harbors a Ty2 element. These shared sequence homologies likely served as a template for break-induced replication (BIR) (**Fig. 2D**), consistent with previously documented Ty-element-mediated BIR events triggered by bleomycin-induced DNA damage in *S. cerevisiae* [22]. We validated this translocation via PCR amplification and Sanger sequencing, confirming that the broken chrXIV end was repaired by BIR using its homologous sequence on chrIV as the template (**Fig. 2E**).

In populations D90_MA_BLM_3 and D90_MA_BLM_5, we observed a recurrent ∼45-kb tandem duplication on chromosome XIII (chrXIII) (**Fig. 2F)**, with paired-end reads confirming the junction bounded by Ty4 elements (**Fig. 2G**). Interestingly, this breakpoint is very close to the native rearrangement of the MA strain that was speculated to experience a historically genomic catastrophic event with multiple chromosome reshuffled [25]. To validate the observation as a true tandem duplication rather than a mapping artifact, we designed outward-facing primers flanking the junction, which yield a PCR product only upon head-to-tail genomic duplication (**Fig. 2H**). Subsequent PCR and Sanger successfully confirmed the Ty4-mediated tandem duplication at this locus (**Fig. 2I**). Interestingly, as noted above, the other two MA populations (D90_MA_BLM_1 and D90_MA_BLM_8) acquired whole-chromosome gain of chrXIII (**Fig. 2A**). The evolutionary convergence of these two distinct mutational routes— tandem duplication and whole-chromosome gain—on the same ∼45-kb interval strongly suggests that upregulation of one or more genes within this region likely drives bleomycin resistance. Further investigation focusing on this locus on the MA background are likely to identify the precise causative gene(s). Together, these findings establish that bleomycin induces structural variation at globally low frequency, but with a highly disproportionate impact on the MA genetic background.

### Base-level mutational landscapes

Although bleomycin is known to elevate single-nucleotide variant (SNV) and insertion-deletion (INDEL) rates in lab yeast strains [22], the extent of this mutagenic susceptibility across diverse backgrounds remains unexplored. The BY lab strain exhibited SNV and INDEL rates of 1.50 × 10^⁻8^ and 1.78 × 10^⁻9^ per base per generation under bleomycin treatment, representing 47-fold and 39-fold higher, respectively, compared to the YPD control (**Fig. 3A-3B, Table S3-S4**). These rates match those previously reported Zheng et al. [22] (Wilcoxon rank-sum test, *P* = 0.65), cross-validating our variant-calling pipeline. However, expansion to five additional genetic backgrounds revealed a striking 10-fold variation in SNV rate, with the MA background exhibiting the lowest rate (1.28 × 10^⁻9^) and the BY background showing uniquely high mutagenic susceptibility (**Fig. 3A**). This finding underscores that mutagenic profiles derived solely from a reference standard lab strain may not generalize across a species. Intriguingly, background-specific mutation rate showed no correlation with either ancestral fitness or fitness gains achieved during the experimental evolution (Pearson correlation, *P* > 0.41). The lack of association implies that drug-induced mutagenesis and adaptive potential are governed by distinct factors. As for the INDEL rate, we detected a unique outlier in one WE population, which is ∼8-fold higher than the other three replicates (**Fig. 3B**). Sequencing of this population and its clonal isolates revealed heterozygous missense and frameshift mutations in *PMS1*, indicating loss of *PMS1* function. Because the *PMS1* endonuclease activity is required for mismatch repair, its functional disruption is expected to drive elevated INDEL rate in this affected population [26,27]. Collectively, these results demonstrate that bleomycin-induced mutagenesis is strongly shaped by genetic background and additional *de novo* mutations in mismatch repair pathways that can dramatically amplify the associated mutational load.

**Figure 3.**
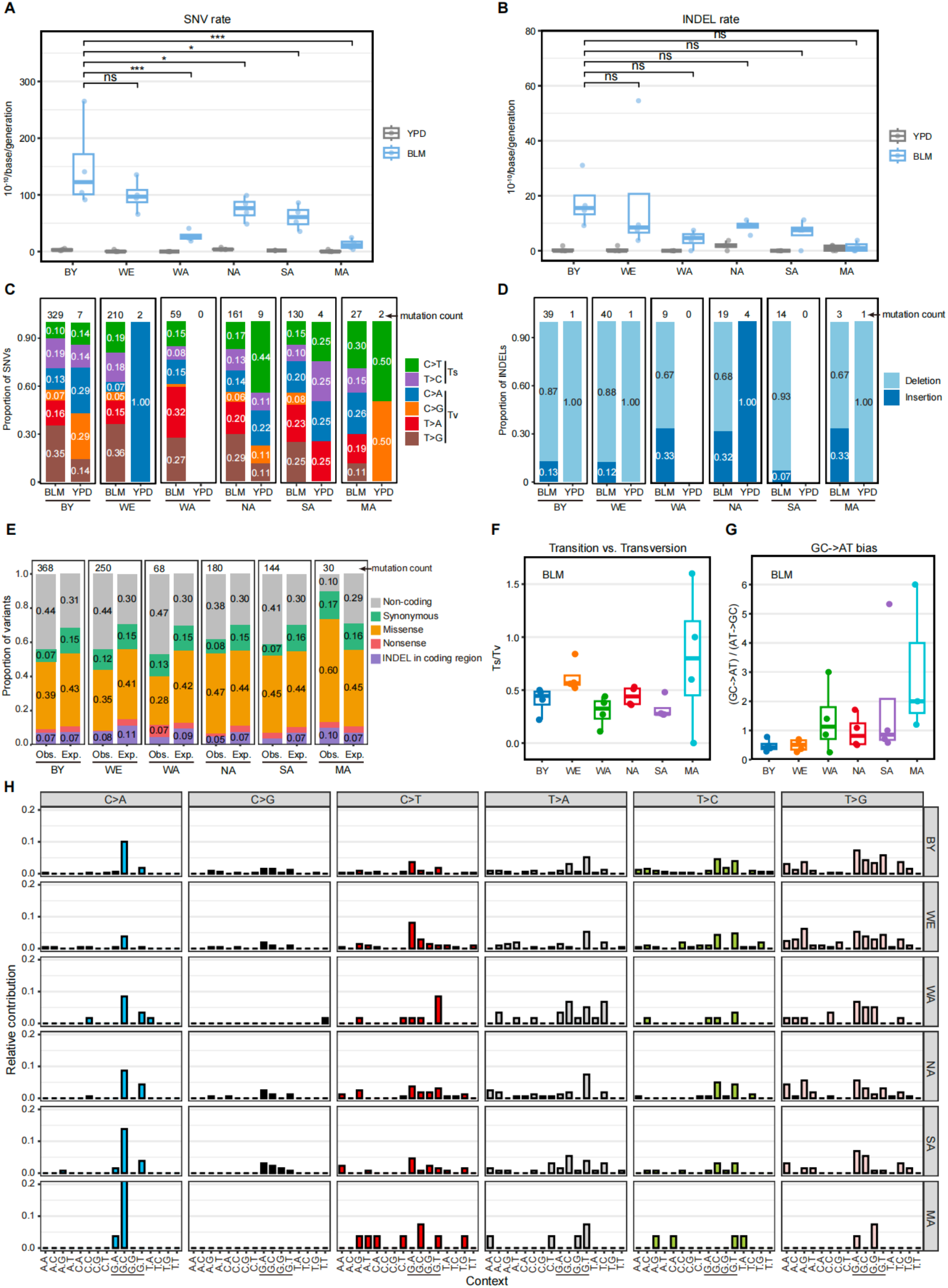
SNV and INDEL features. **(A)** SNV and **(B)** INDEL rates across genetic backgrounds in YPD and bleomycin conditions. Box plots represent the median, interquartile range, and min-to-max range, overlaid with individual data points. Rates are scaled by 10^−10^ per base per generation. Statistical significance among bleomycin-treated groups was assessed using Dunnett’s post-hoc test. Asterisks indicate significant differences relative to BY: *p < 0.05, p < 0.01, ***p < 0.001; ns, not significant. Distribution of **(C)** SNV and **(D)** INDEL spectra across genetic backgrounds in YPD and bleomycin conditions. Bars represent the proportion of each substitution type. Mutation counts were aggregated across four independently evolved populations per background. Differences in SNV mutational spectra among bleomycin-treated groups were assessed using a chi-square test based on substitution counts (Chi-Square test, χ² = 59.03, df = 25, p = 0.0001). **(E)** Comparison of observed and expected mutation consequence spectra across genetic backgrounds. Obs., observed mutation spectrum; Exp., background-specific neutral expectation. **(F)** Ts/Tv and **(G)** GC-to-AT bias across genetic backgrounds in bleomycin. **(H)** Mutational profiles based on the relative incidences of SNV within a trinucleotide context across genetic backgrounds in bleomycin.

Regarding SNV spectrum, under the standard YPD condition, the spontaneous SNV typically display two characteristic patterns: (1) a high transition-to-transversion ratio (Ts/Tv, e.g., ∼0.93-0.95) driven by frequent C-to-T transition, and (2) a pronounced GC-to-AT mutational bias arising from elevated rate of C-to-T transition and C-to-A transversion [11,28]. Bleomycin exposure remarkably reshaped the spectra. It increased the proportion of T-to-G and T-to-A transversion while reducing the proportion of C-to-T transition, thereby markedly resulting in a significantly lower Ts/Tv ratio (**Fig. 3C and 3F**). Together with the decreased proportion of C-to-A changes, bleomycin shifted the mutational spectrum away from the canonical spontaneous GC-to-AT bias (**Fig. 3G**).

Beyond these shared signatures, bleomycin-induced SNV mutational spectra differed significantly among genetic backgrounds (**Fig. 3C,** *P* = 0.0001). When evaluating the genome-wide distribution of mutational consequences against neutral expectations, we observed significant variation among the six backgrounds (**Fig. 3E**, Pearson’s χ² test of independence with Monte Carlo-simulated *P* values based on 100,000 permutations; *P* = 0.0034). Specifically, five backgrounds (BY, WE, WA, NA, and SA) individually deviated from their own background-specific neutral expectations (**Fig. 3E**, Monte Carlo-simulated Pearson’s χ² goodness-of-fit test, BH-adjusted *P* < 0.05). Furthermore, in stark contrast to the YPD control condition where neighboring bases of *de novo* SNVs are predominantly enriched for <u>C</u>C<u>G</u> and <u>T</u>C<u>G</u> context [28], bleomycin-induced SNVs displayed a strong preference for the guanine (G) as the immediate 5′ neighbor (**Fig. 3H**). This is consistent with the well-established susceptibility of the 5′-G-pyrimidine-3′ motif to bleomycin-induced DNA lesions. Notably, a robust <u>G</u>C<u>C</u> signature persisted across all six genetic backgrounds (**Fig. 3H**), indicating that this sequence preference is a conserved, background-independent hallmark of bleomycin mutagenesis.

Regarding INDELs, the single-base-pair deletion represented the predominant class (mean = 81.5%), which is the characteristic of error-prone translesion synthesis (TLS) bypassing bleomycin-induced apurinic/apyrimidinic (AP) sites. However, the exact proportion of the 1-bp deletions varied considerably across genetic backgrounds (66.7% ∼ 92.9%) (**Fig. 3D**), indicating that the efficiency or regulation of TLS pathway may differ across yeast strains of distinct origins. In addition, approximately half of the detected INDELs (62/124) occurred within genic regions, closely matching the proportion observed in the YPD control condition (53%) [28]. This indicates that while bleomycin exposure accelerates INDEL accumulation, it does not fundamentally alter their genomic distribution.

Taken together, these results reveal that bleomycin mutagenesis comprises a conserved biochemical signature, reflecting the drug’s shared mechanism of DNA damage. Nevertheless, we also uncover a critical layer of background-dependent variation in both the mutation rate and mutational spectrum. Therefore, the ancestral genetic background acts as a potent modifier that scales the magnitude of bleomycin-induced genome instability without altering its fundamental molecular character.

### Driver mutations for bleomycin resistance

We next asked whether bleomycin-induced mutagenesis gives rise to mutations that drive resistance. Across the six genetic backgrounds, we identified recurrent mutations in 11 genes (**Fig. 4A**). Among them, *PTK2* is a well-established regulator of bleomycin resistance [20]. It encodes a serine/threonine protein kinase that regulates ion transport across the plasma membrane and its loss-of-function impairs drug uptake. This mechanism aligns well with our finding that nearly all identified *PTK2* mutations in bleomycin-evolved BY, WE and WA populations were homozygous frameshift mutations that interrupt its kinase function. We also recaptured mutations in *SKY1*, another known driver of bleomycin resistance, in evolved WE and MA populations. *SKY1* encodes a kinase regulating cation homeostasis through membrane transporters, and its deficiency is known to confer bleomycin resistance [20,22]. In addition, we identified *PMA1* as a novel candidate for driving bleomycin resistance. Strikingly, it emerged as the most frequently mutated gene across all the backgrounds. *PMA1* encodes the plasma membrane P2-type H^+^-ATPase responsible for regulating cytoplasmic pH and plasma membrane potential. But its functional link to bleomycin resistance, to our knowledge, was previously unknown.

**Figure 4.**
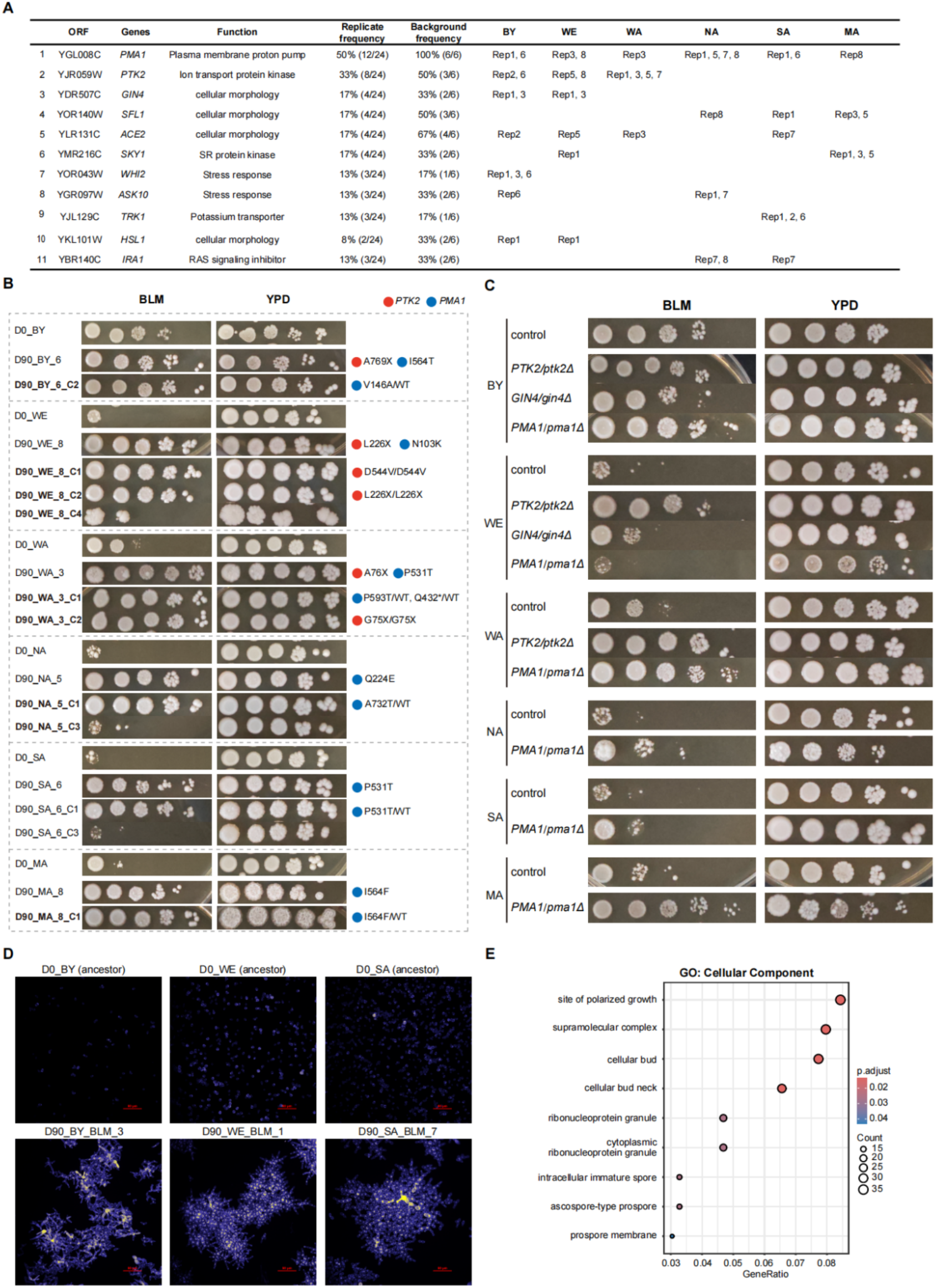
Driver mutations for bleomycin resistance. **(A)** Recurrent mutated genes across evolved populations. **(B)** Phenotypic validation of populations and clones with *pma1* and *ptk2* mutations. **(C)** Phenotypic validation of constructed clones with *pma1*, *ptk2* and *gin4* deletions. **(D)** Cellular elongation and aggregation in bleomycin-evolved populations. **(E)** GO (cellular component) analysis of bleomycin-specific mutations with moderate or high effects.

To determine whether the recurrent mutations confer resistance, we validated the causal role of the top two frequently occurring mutations, *PMA1* and *PTK2,* by genotyping and phenotyping the clones isolated from the evolved populations carrying these mutations (**Fig. 4B, Table S5-S7**). To avoid confounding effects from co-occurring mutations, we restricted our primary analysis to clones carrying only one candidate mutated gene (**Figure 4B**, see the full phenotyping dataset in **Fig. S4**). Clones carrying *PMA1* or *PTK2* mutations exhibited near-complete resistance to bleomycin in stark contrast with their wildtype parents. Notably, *PMA1* and *PTK2* mutations are mutually exclusive across our sequenced clones, supporting their roles as major evolutionary drivers. To further validate the gene function directly, we generated targeted gene deletion in their respective ancestral backgrounds where the corresponding mutations were identified. Given *PMA1* is an essential gene, we constructed heterozygous deletion mutants (*PMA1/pma1Δ*). Heterozygous deletion of *PMA1* leads to bleomycin resistance although the magnitude of the resistance is background dependent (**Fig. 4C**). Furthermore, the heterozygous *PMA1* deletion in WE, NA and MA backgrounds exhibits growth defects in YPD condition, suggesting a background-dependent haploinsufficiency of this essential proton pump. The absence of such phenotypes in BY, WA and SA backgrounds underscore the substantial influence of genetic background on gene essentiality. *PTK2* deletion in BY, WE and WA backgrounds, where we identified this mutation in their respective evolved populations, all confer strong bleomycin resistance (**Fig. 4C**). Together, these results establish *PMA1* and *PTK2* loss-of-function as the major drivers of bleomycin resistance during adaptive evolution.

In addition to these primary drivers, the remaining mutated genes have not previously been linked to bleomycin resistance and may represent hitchhiker mutations or contribute additively (**Fig. 4A**). *GIN4*, encodes a protein kinase involved in bud growth and septin ring assembly. Clones with *GIN4* mutation confer mild resistance though still significantly better than the wildtype parent (**Fig. S4)** and *GIN4* deletion produced moderate resistance (**Fig. 4C**). Interestingly, besides *GIN4,* another three mutated genes, *SFL1, ACE2* and *HSL1,* function in regulating cellular morphology and aggregation, are distributed sporadically across different genetic backgrounds. Consistent with their functions, we observed a significantly elevated proportion of populations displaying elongated buds and cell-separation-deficient phenotypes under bleomycin condition whereas no such complex phenotype was observed in YPD control condition (39.6% *vs.* 0%; **Fig. 4D, Table S8**). Moreover, gene ontology (GO) analysis of all mutations with high or moderate effect in bleomycin further revealed enrichment for polarized growth, cellular bud, and bud neck components (**Fig. 4E, Fig. S5**).

Together, our results validated previously documented bleomycin-resistant drivers such as *PTK2* and *SKY1*, while establishing *PMA1* as a novel and background-independent driver. Furthermore, our characterization of other recurrent mutations revealed several secondary modulators associated with cellular morphology, whose precise mechanistic contributions to drug tolerance remain to be fully defined (see Discussion).

### Mechanisms of *PMA1* and *PTK2* mutations to drive bleomycin resistance

The convergence of *PMA1* and *PTK2* mutations across nearly all genetic backgrounds suggested a shared mechanistic basis for bleomycin resistance. Pma1p is the primary plasma membrane H^+^-ATPase that exports protons to establish the plasma membrane potential [29] while Ptk2p is a known positive regulator of Pma1p activity [30]. Given this functional connection, we hypothesized that *PMA1* and *PTK2* mutations confer resistance by disrupting Pma1p ATPase activity, thereby depolarizing the plasma membrane and limiting entry of the positively charged bleomycin molecule. Supporting this hypothesis, all *PMA1* mutations mapped within the P-type ATPase functional domain, suggesting a direct impairment of its ATPase activity (**Fig. 5A)**. Similarly, almost all the identified *PTK2* mutations were loss-of-function variants (e.g., frameshift) (**Fig. 5B)** and *ptk2Δ* cells are known to have significantly reduced Pma1p activity [31]. Together, these observations support a model in which both *PMA1* and *PTK2* mutations converge on the impairment of Pma1p function.

**Figure 5.**
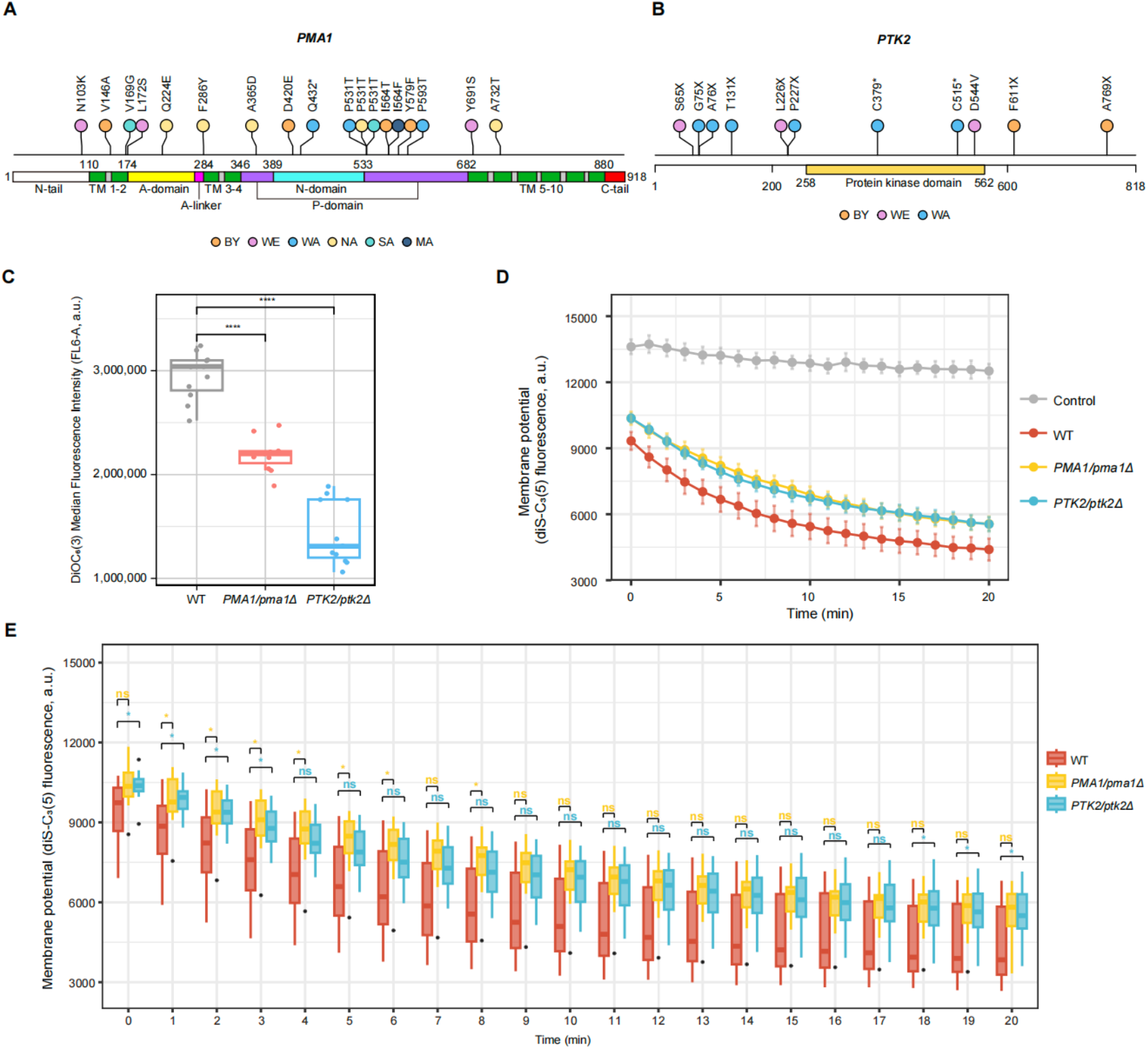
Mechanism of *PMA1* and *PTK2* conferring bleomycin resistance. **(A)** Mutation positions and their corresponding functional domains of *PMA1* [34]. **(B)** Mutation positions and their corresponding functional domains of *PTK2*. **(C)** Plasma membrane potential measurement using DiOC_6_(3) by flow cytometry. **(D-E)** Plasma membrane potential measurement using diS-C_3_(5) by plate reader.

To test this hypothesis directly, we measured plasma membrane potential using two independent complementary methods. First, we used DiOC_6_(3), a cationic cyanine lipophilic dye that passively diffuses into the cells relying on the plasma membrane potential [32]. Thus, its cellular fluorescence intensity is a readout of plasma membrane potential. Flow cytometry revealed that the DiOC_6_(3) fluorescence intensity significantly reduced in *pma1Δ* and *ptk2Δ* constructs compared to their wildtype control (**Fig. 5C, Fig. S6**), indicating membrane depolarizations. Because DiOC_6_(3) signal can be confounded by the mitochondria membrane potential, we corroborated this result using a second mechanistically distinct probe, diS-C_3_(5) [33]. This dye enters polarized cells and undergoes self-quenching at high cellular concentrations, whereas cell depolarization impairs its accumulation and yields higher fluorescence with slower quenching kinetics. Consistent with our initial readout, we observed significantly elevated fluorescence and delayed quenching kinetics in *pma1*Δ and *ptk2*Δ constructs compared to wild-type (**Fig. 5D-5E**), again confirming membrane depolarizations. Together, by combining functional prediction with two independent and mechanistically distinct fluorescence-based assays, we validate our hypothesis that *PMA1* and *PTK2* mutations disrupt the plasma membrane potential by impairing Pma1p H+-ATPase activity. This attenuated membrane potential provides a unifying, background-independent mechanism of bleomycin resistance: by reducing the electrochemical driving force for uptake of the positively charged drug, it limits intracellular bleomycin entry into the cell irrespective of the genetic background (**Fig. 6**).

**Figure 6.**
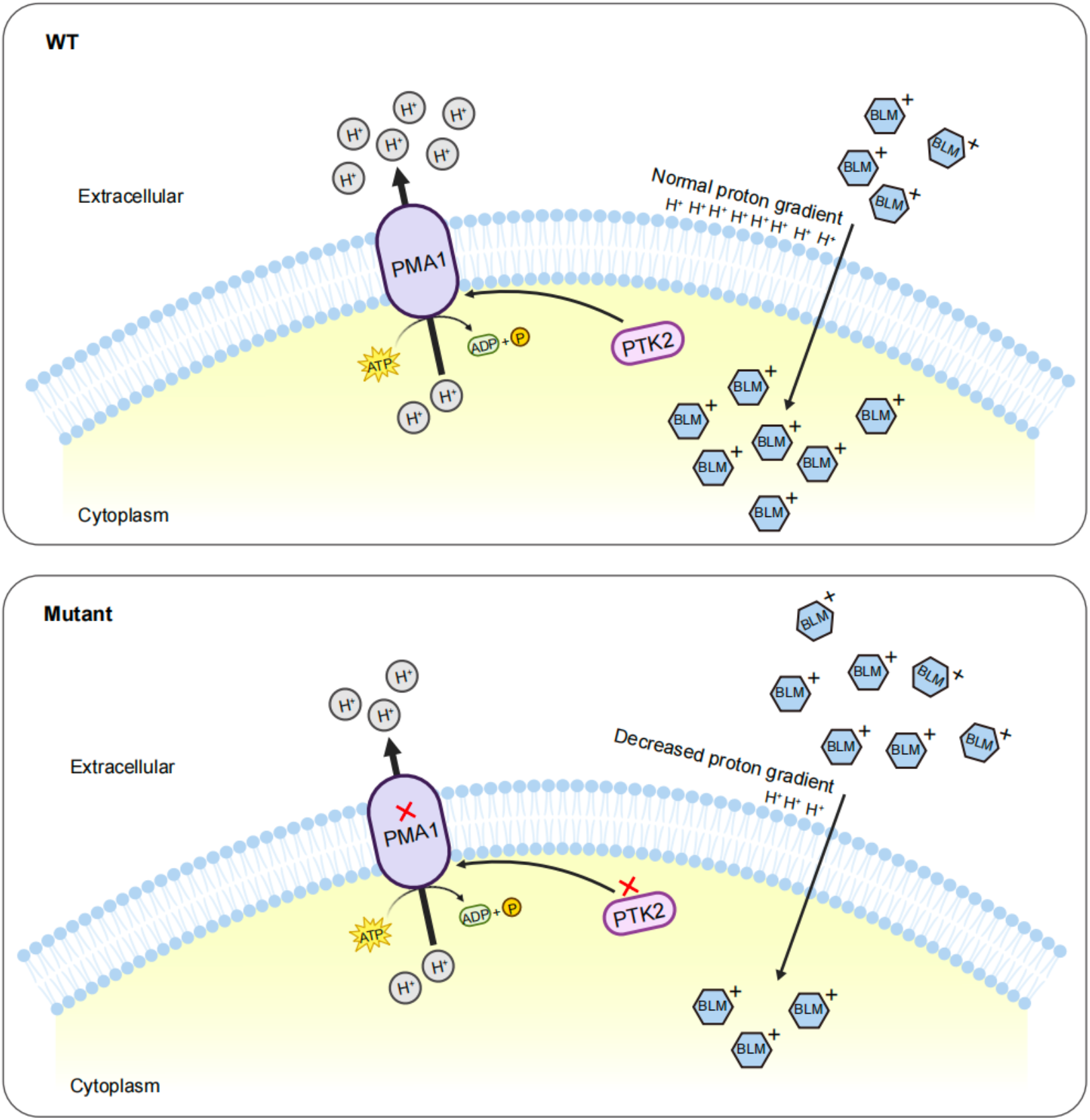
Working model of *PMA1* and *PTK2* mediating bleomycin resistance.

## Discussion

Through the experimental evolution of six genetically and geographically distinct *S. cerevisiae* strains under bleomycin selection, we demonstrated that while resistance arises repeatably, the underlying paths are only partially predictable from the drug’s known mechanism of action. We found that genetic background profoundly shapes the magnitude and reproducibility of adaptation, the resulting genomic footprints, and the mutational raw material accessible to selection. Nonetheless, our study uncovered a highly convergent, background-independent biochemical mechanism of resistance mediated by *PMA1* and *PTK2* mutations, illustrating how universal physiological constraints can guide evolutionary outcomes despite broad genomic divergence.

The phenotypic variance among backgrounds, together with the inverse relationship between ancestral fitness and fitness gain, fits the model of diminishing-returns epistasis in which the same mutational target confers less benefit as a genotype approaches a local fitness optimum [35]. However, diminishing returns alone may not fully explain why backgrounds differ in their adaptive ceilings rather than merely their rates of adaptation. A more intriguing possibility is that pre-existing variants segregating within each background modulate the fitness effects of newly arising resistance mutations—a form of background-by-mutation epistasis that alters beneficial or deleterious effects of a mutation depending on the genomic context where it emerges. If so, the bleomycin sensitive MA replicates may not simply have failed to acquire a driver mutation in 90 days, but may occupy a genetic background in which the mutation-selection balance for these alleles is shifted, either because their fitness benefit is attenuated or the pleiotropic cost is amplified. This would echo growing evidence that epistasis with genetic background, rather than mutation supply *per se*, frequently limits evolutionary parallelism [14].

While traditional studies on adaptive evolution have frequently relied on the standard lab strains like BY to dissect the mechanics of drug resistance, our findings sound a cautionary note against this single-background paradigm. Notably, we detected a striking 10-fold variance in bleomycin-induced SNV rates across six genetic backgrounds, revealing that the canonical lab strain is exceptionally prone to mutagenicity. That the BY lab strain sits at the extreme high bound of this spectrum is highly telling: generations of domestication and lab passage may have eroded stress-responsive mutational buffering retained in wild isolates, rendering BY a mutagenic outlier rather than a representative baseline. Such high level of bleomycin mutagenesis of BY also introduces significant caveats when using bleomycin as a selective marker for genome editing, as it may inadvertently bring confounding off-target mutations during screening. Alongside recent works showing that loss of heterozygosity (LOH) rates varied widely across different yeast strains [36,37], our work underscores how an overreliance on a single lab strain can potentially distort our view of drug-induced mutagenesis.

Bleomycin induced aneuploidy and copy number associated SV were rare but enriched in specific backgrounds, like MA and SA. While it remains to be determined whether these events directly drive adaptation, copy number alterations that modulate gene expression are well-known to mediate phenotypic adaptation [38,39]. The genomic events observed on chromosome XIII in the MA strain offer a fascinating example of evolutionary convergence at the locus level achieved through completely different structural mechanisms. Specifically, two independent MA populations achieved copy number gains via whole-chromosome XIII aneuploidy while two other MA populations acquired ∼45-kb tandem duplication on the same chrXIII. This locus-specific convergence implies that the upregulation of one or more genes within this 45-kb interval provides a potent, background-dependent fitness advantage under bleomycin selection, as such high recurrence at the same locus across independent populations is hard to attribute to drift alone.

The high recurrence of mutations in *PMA1* and *PTK2* across multiple genetic backgrounds points to a highly constrained, shared mechanism of resistance. Pmalp is the primary proton pump regulating cytoplasmic pH and plasma membrane potential, while Ptk2p acts as its positive regulator [30]. The loss-of-function mutations in *PTK2* and targeted mutations within the P-type ATPase domain of *PMA1* systematically disrupt this system. By decreasing the plasma membrane potential, the cells effectively build a biophysical shield that blocks the influx of positively charged bleomycin molecules. Intriguingly, while the mechanism of resistance is universal, its evolutionary cost is not. Heterozygous deletions of *PMA1* triggered notable growth defects and haploinsufficiency in the WE, NA, and MA backgrounds, yet left the BY, WA, and SA backgrounds completely unaffected. This variation in fitness costs highlights how pre-existing epistatic networks dictate the long-term viability of specific resistance mutations. Furthermore, while *ptk2* mutation has been identified to generate bleomycin resistance by independent studies [20,22], to our knowledge this study provides the first validation that *pma1* mutation directly confers bleomycin resistance independent of genetic backgrounds.

Our study also identified an enrichment of mutations in genes regulating cellular morphology and polarized growth, including *GIN4*, *SFL1*, *ACE2*, and *HSL1*. These mutations correlated with a striking shift toward elongated buds and multicellular-like aggregation phenotypes unique to the bleomycin condition. While *GIN4* mutations were validated to confer moderate bleomycin resistance directly, the effects of other mutations remain to be understood. It also remains an open question whether these macro-structural cellular alterations serve as an active defense mechanism [40,41] - perhaps by reducing the relative cell surface area, altering cell wall permeability, or whether they represent a tolerable downstream consequence of altered cell-cycle kinetics under chronic DNA damage stress.

By scaling experimental evolution from a single lab background to a diverse panel of genetic backgrounds, this study mirrors the complex clinical realities of human oncology and infectious disease. Patients with distinct genetic backgrounds exhibit highly variable toxicities, drug resistance kinetics, and therapeutic outcomes when treated with standard chemotherapies like bleomycin. Our study based on the yeast model demonstrated that genetic background fundamentally modulates drug tolerance and mutational susceptibility. Elucidating the interplay between background-specific constraints and universal biophysical vulnerabilities (such as membrane potential modulation) provides a dual framework for designing more robust, individual-specific therapeutic strategies to circumvent the evolution of drug resistance.

## Materials and Methods

### Experimental Evolution

Six diploid homozygous *Saccharomyces cerevisiae* strains, derived from backgrounds representing diverse ecological niches, BY4743 (lab strain, BY), DBVPG6765 (Wine/European, WE), DBVPG6044 (West African, WA), YPS128 (North American, NA), Y12 (Sake, SA), and UWOPS03-461.4 (Malaysia, MA), were utilized in this study. For each background, a single colony was isolated to serve as the ancestor (Day 0). Experimental evolution was conducted in parallel under two environments: the selective condition in YPD broth media (Sangon Biotech, Cat. No. A507022-0250) supplemented with 12.5 μg/mL bleomycin (MCE, Cat. No. HY-17565) and the control condition in drug-free YPD.

Eight independent replicate lines were established per background for each condition, yielding a total of 48 lines per environment (6 backgrounds × 8 replicates). Serial transfers were performed every 48 hours by inoculating 5 μL of thoroughly mixed culture into 150 μL of fresh medium (with or without bleomycin), followed by static incubation at 30 °C. Each transfer cycle corresponded to approximately 5 generations (log_2_ [(150+5)/5] ≈ 4.95). Over a 90-day period, a total of 45 serial transfers were completed, accumulating to an estimated 225 generations per replicate line. To facilitate downstream phenotypic and genotypic tracking, culture aliquots were cryopreserved at Days 0, 2, 4, 8, 16, 20, and every 10 days thereafter.

### Whole genome sequencing

Genomic DNA (gDNA) was extracted from six ancestral strains, as well as four replicate populations per genetic background and the isolated clones at the final time point (Day 90). To ensure representative population sampling, 10 μL of each cryopreserved culture was inoculated into 100 μL of YPD medium and pre-cultured overnight at 30 °C. The pre-culture was subsequently transferred into 5 mL of the respective evolution medium and incubated overnight at 30 °C with shaking at 220 rpm.

The gDNA extraction was performed using the MasterPure Yeast DNA Purification Kit (Lucigen, Cat. No. MPY80200) with several modifications. Briefly, cell wall digestion was carried out by treating samples with 20 μL of lyticase (200 units/mL; Sigma-Aldrich, Cat. No. L4025-50KU) at 37 °C for 50 min. To remove RNA, 20 μL of RNase A (10 mg/mL; MP Biomedicals, Cat. No. 101076) was added, followed by incubation at room temperature for 90 min. During both enzymatic digestion steps, the mixtures were vortexed every 10 minutes to maintain homogeneity. Finally, paired-end sequencing (2 × 150 bp) was performed on the DNBSEQ platform (MGI Tech Co., Ltd., Shenzhen, China), yielding mean sequencing coverage is 173× (**Table S2**).

### Phenotypic analysis

Phenotypic profiling was performed using both spotting assays and growth curve analysis. Cells recovered from cryopreserved stocks were subjected to ten-fold serial dilutions (10⁻¹ to 10⁻⁴). Subsequently, 3.5 μL of each dilution were spotted onto YPD agar plates (Sangon Biotech, Cat. No. A507023-0250) in the presence or absence of bleomycin. Plates were incubated at 30 °C for 72 hours prior to photography.

For kinetic growth analysis, the recovered cultures were diluted ten-fold, and 2 μL of the diluted suspension was inoculated into 150 μL of YPD medium with or without drug in a 96-well plate. Growth dynamics was monitored using an Epoch 2 microplate reader (BioTek), and generation time, lag time and yield were calculated using PRECOG software [42]. All downstream data visualization and statistical plotting were executed in R (version 4.5.1).

### Genomic data analysis

Genomic data from all ancestral strains, evolved populations, and isolated clones were processed using a unified pipeline. Reference genome sequences for BY, WE, WA, NA, SA, and MA backgrounds were retrieved from the repository via the link https://yjx1217.github.io/Yeast_PacBio_2016/data/ [25]. Read mapping and variant calling, including SNVs, INDEL, SV, and CNV, were executed via the Varathon pipeline (https://github.com/yjx1217/Varathon). Specifically, SNV and INDEL were identified using FreeBayes [43] and GATK4 [44] as variant callers, while SV was resolved using Manta [45] and Delly [46].

For SNV and INDEL filtering, quality thresholds were applied using BCFtools (v1.14) [47]. GATK-derived variants were retained based on the following criteria: FS < 60, MQ > 40, MQRankSum > −3, QD > 3, ReadPosRankSum > −2, and SOR < 3. FreeBayes-derived calls were filtered using the thresholds of EPP > 3, SAP > 3, and SRP > 3. The intersection of filtered calls from both tools was defined as the consensus variant set, which was subsequently merged across samples within each genetic background. Functional annotation was performed using the Ensembl Variant Effect Predictor (VEP) (v114.2) [48]. Variants situated within repetitive regions, telomeres, and subtelomeres were excluded. To eliminate low-coverage artifacts, candidate sites were required to have a total read depth ≥ 10 and an alternate allele depth ≥ 3. Baseline variants pre-existing in the ancestral genomes (Day 0) were removed. All high-confidence SNVs and INDELs passing these filters were visually validated using the Integrative Genomics Viewer (IGV, v2.19.5) [49]. For downstream analysis, systematic gene IDs for mutated loci were converted to standard gene symbols using the R package *org.Sc.sgd.db* (v3.21.0). To characterize the mutational landscape, 96-trinucleotide mutation profiles were generated with the *MutationalPatterns* R package (v3.18.0). For Gene Ontology (GO) enrichment analysis, variants predicted to have moderate or high functional impacts were retained and deduplicated at the gene level. Enrichment and subsequent visualizations were executed using the *clusterProfiler* (v4.16.0) and *enrichplot* (v1.28.4) R packages. Finally, lollipop plots of *PMA1* and *PTK2* were created using the *trackViewer* R package (v1.44.0).

SV candidates identified by Manta and Delly were filtered independently, requiring a minimum of 5 supporting reads. Ancestral SV background was subtracted by detecting genomic coordinate overlaps using the R package *GenomicRanges* (v1.60.0), with a maximum gap tolerance of 100 bp. An SV was discarded if it overlapped spatially and shared an identical variant type (SVTYPE) with the ancestral baseline. A comprehensive union set of SVs was then constructed for each sample by combining the filtered datasets from both callers, retaining both caller-specific and mutually supported variants. For overlapping variants called by both tools, coordinates and annotations from Manta were preferentially retained. All candidate SVs were subjected to visual inspection in IGV to eliminate potential false positives.

### Calculation of expected mutation spectra

Expected mutation spectra were estimated separately for each genetic background based on genome architecture and the theoretical distribution of mutation consequences. For each background, the proportions of coding and non-coding regions were calculated from the corresponding genome annotation (GFF) file. Based on the total numbers of observed SNVs and INDELs in each background, expected mutation counts were calculated according to these genomic proportions. Specifically, the expected number of non-coding mutations was calculated as the sum of expected non-coding SNVs and expected non-coding INDELs. The expected number of INDELs occurring within protein-coding regions was calculated based on the proportion of coding sequences in each genome. For SNVs occurring within coding regions, the expected proportions of synonymous, missense, and nonsense mutations were estimated according to the theoretical distribution derived from the genetic code (24.6%, 69.6%, and 5.8%, respectively), resulting in the expected numbers of synonymous, missense, and nonsense mutations. The resulting expected mutation spectra were compared with the observed mutation consequence spectra for each genetic background. Differences between observed and expected mutation spectra were assessed using Monte Carlo-simulated Pearson’s χ² goodness-of-fit tests with 100,000 replicates. P values were adjusted for multiple testing using the Benjamini-Hochberg (BH) method.

### Structural variant validation

We validated the SVs as shown in Figure 2. Per-base mapping depth outputs from Varathon were utilized to calculate median sequencing depths across non-overlapping 10-kb windows. Candidate genomic regions representing copy number gains or losses were visually inspected at the single-nucleotide breakpoint level using the IGV. To experimentally verify the breakpoints, locus-specific primers were designed based on the Saccharomyces Genome

Database (SGD) [50] and synthesized by Tsingke Biotechnology (Beijing, China) (**Table S9**). PCR amplifications were performed using 2 × Phanta Flash Master Mix (Vazyme, P510) according to the manufacturer’s instruction. The resulting amplicons were resolved by 1% agarose gel electrophoresis and subsequently submitted to Tsingke Biotechnology Co., Ltd. for Sanger sequencing. Sequencing chromatograms were inspected and sequence alignments with the corresponding reference sequences were performed using SnapGene software (v.8.0; www.snapgene.com).

### Target gene deletion and yeast transformation

Target gene deletion primers were designed using the online tool GetPrimes [51] (https://www.evomicslab.org/app/getprimers/) based on the respective genetic backgrounds. The deletion primers contained a 5′ 40-bp homology arm complementary to the sequences immediately flanking the target open reading frame (ORF), enabling homologous recombination with the *URA3* selectable marker. The linear *URA3* knockout cassette was amplified via PCR using the *URA3*-containing pYES plasmid as the template. The PCR product is ready for transformation. Verification primers situated further upstream and downstream of the target locus were also designed and synthesized (**Table S9**).

For transformation, wild-type yeast strains were revived and streaked for single colonies on YPD agar plates. A single colony was inoculated into 5 mL of YPD liquid medium and cultured overnight at 30 °C with shaking at 220 rpm. For each transformation, a 250-μL aliquot of the overnight culture was transferred into 5 mL of fresh YPD medium and grown for an additional 4–5 h. A positive control group of direct plasmid transformation was prepared in parallel for each genetic background.

Cells were then harvested by centrifugation at 2,500 rpm for 5 min, and the supernatant was discarded. The pellet was resuspended in 700 μL of sterile water, transferred to a 1.5-mL microcentrifuge tube, and centrifuged at 10,000 rpm for 1.5 min. The cells were then gently resuspended in 500 μL of 0.1 M lithium acetate (LiAc, Sigma-Aldrich, Cat. No. L4158) and pelleted again. After thoroughly removing the supernatant, the transformation components were sequentially added to the cell pellet: 240 μL of 50% (w/v) PEG 3350 (Sigma-Aldrich, Cat. No. P4338), 36 μL of 1 M LiAc, 80 μL of the PCR product (or ∼1 μg of intact plasmid adjusted to 80 μL with sterile water for the positive control), and 10 μL of single-stranded salmon sperm DNA (pre-boiled at 99 °C for 5 min and chilled immediately on ice). The mixture was vortexed thoroughly, incubated at 30 °C for 30 min, and subjected to heat shock at 42 °C for 30 min. Following heat shock, cells were pelleted at 8,000 rpm for 2 min, washed with 900 μL of sterile water, and resuspended in 100 μL of sterile water. A 50-μL aliquot of the cell suspension was plated onto synthetic complete agar plates deficient for uracil (SC–Ura), containing 0.68% YNB (Sigma-Aldrich, Cat. No. Y0626), 2% glucose (Sigma-Aldrich, Cat. No. G8270), 0.08%

Ura-dropout amino acid mix (MP Biomedicals, Cat. No. 4511-222), and 2% agar (Sangon Biotech, Cat. No. A505255-0250), with the pH adjusted to 6.0–6.5 using NaOH. Prototrophic transformants were selected after incubation at 30 °C. Gene deletion (replaced by *URA3* selective marker) was verified by colony PCR using the verification primers, followed by 1% agarose gel electrophoresis. Confirmed knockout strains were preserved in 25% glycerol at −80 °C.

### Calcofluor White staining of cell morphology

Cell morphology was assessed by Calcofluor White (Sigma, Cat. No. 18909) staining following a previously described protocol [52]. Briefly, 20 µL of culture was pelleted by centrifugation, stained with 25 µL of 1:5 diluted Calcofluor White in the dark for 5 min, washed once with 25 µL ddH_2_O, and resuspended in 5 µL ddH_2_O. Stained cells were imaged using a NIKON Eclipse Ni-U fluorescence microscope.

### Membrane potential measurement

#### DiOC₆(3) Staining and Flow Cytometry Analysis

To evaluate the plasma membrane potential, staining with the cationic cyanine lipophilic probe 3,3’-dihexyloxacarbocyanine iodide [DiOC_6_(3); Solarbio, Cat. No. D3850] was performed according to previously established method [32]. Yeast strains (wild-type, *ptk2Δ*, and *pma1Δ*) were pre-cultured overnight at 30 °C. Subsequently, 250 μL of each culture was inoculated into 5 mL of YPD medium and incubated at 30 °C with shaking at 220 rpm for 5 h to reach the logarithmic growth phase.

Cells were harvested by centrifugation at 2,500 rpm for 4 min, and the supernatant was discarded. The cell pellets were washed with 1 mL of sterile water, transferred to 1.5-mL microcentrifuge tubes, and centrifuged again at 3,000 rpm for 2 min. The washed cells were resuspended in 1 mL of sterile water, and their optical densities at 600 nm (OD_600_) were determined. The cell suspensions were then standardized to an OD_600_ of 0.2 in a final volume of 1 mL. For each sample, an unstained aliquot was maintained as a negative control. To the remaining tubes (n=11 for each strain), a 23-μL aliquot of the working dye solution was added to achieve a final dye concentration of 0.4 µM. The mixtures were incubated at 37 °C for 10 min in the dark using an aluminum foil-covered metal bath. Following incubation, cells were collected by centrifugation at 1,800 rpm for 5 min, and the pellets were sequentially resuspended in 1 mL of sterile water immediately prior to analysis.

Flow cytometry was performed on a CytoFLEX flow cytometer equipped with a 488 nm argon-ion laser as the excitation source. The green fluorescence was detected using the 530 ± 40 nm channel. For each sample, a total of 30,000 events were acquired with forward scatter (FSC) and side scatter (SSC) recorded simultaneously.

#### diS-C3(5) Fluorescence Self-Quenching Assay

To complement the flow cytometry findings, the membrane potential was further validated via a microplate assay using the 3,3’-dipropylthiodicarbocyanine iodide probes (diS-C3(5); Macklin, Cat. No. D864435) based on its concentration-dependent self-quenching properties upon intracellular accumulation [33]. The diS-C3(5) stock was prepared at 0.1 mM in ethanol and stored at -20 °C in the dark. The assay buffer consisted of 10 mM Na_2_HPO_4_ (Solarbio, Cat. No. D7290) and is adjusted to pH 6.0 with HCl.

Yeast strains were cultured and harvested under identical conditions to those described above. After washing with sterile water, the cell pellets were resuspended in the assay buffer, and the cell density was adjusted to an OD_600_ of 0.05. A 150-μL aliquot of the adjusted cell suspension was transferred into each well of a 96-well plate. Staining was initiated by adding 6 μL of the diS-C3(5) working solution to each well to obtain a final concentration of 4 µM. Fluorescence intensity was monitored immediately using a BioTek Synergy H1 multimode microplate reader, with excitation and emission wavelengths set at 600 nm and 675 nm, respectively. Kinetic readings were recorded every minute over a 20-min duration (n=10 for each strain). Unstained cells (cells in buffer only, n=3) and blank controls (buffer with dye only, n=3) were processed in parallel.

## Data availability

The raw sequence data reported in this paper have been deposited in the Genome Sequence Archive [53] in National Genomics Data Center [54], China National Center for Bioinformation / Beijing Institute of Genomics, Chinese Academy of Sciences (GSA: CRA046458) that are publicly accessible at https://ngdc.cncb.ac.cn/gsa/s/27w5Eh7m. The project ID is PRJCA069003

## Acknowledgements

This work is supported by the Young Talents Program of Sun Yat-sen University Cancer Center (YTP-SYSUCC-0040 to J.L., YTP-SYSUCC-0042 to J.-X. Y.), the National Natural Science Foundation of China (32470663 to J.-X. Y., U25C2027 to L. Z.), and the Key Research and Development Program of Ningxia Hui Autonomous Region (2026BBF01002 to J.-X. Y.).

