## Supplementary Figures for "Genetic background diversifies mutational paths but converges on membrane depolarization for bleomycin resistance in yeast"

**Figure S1**

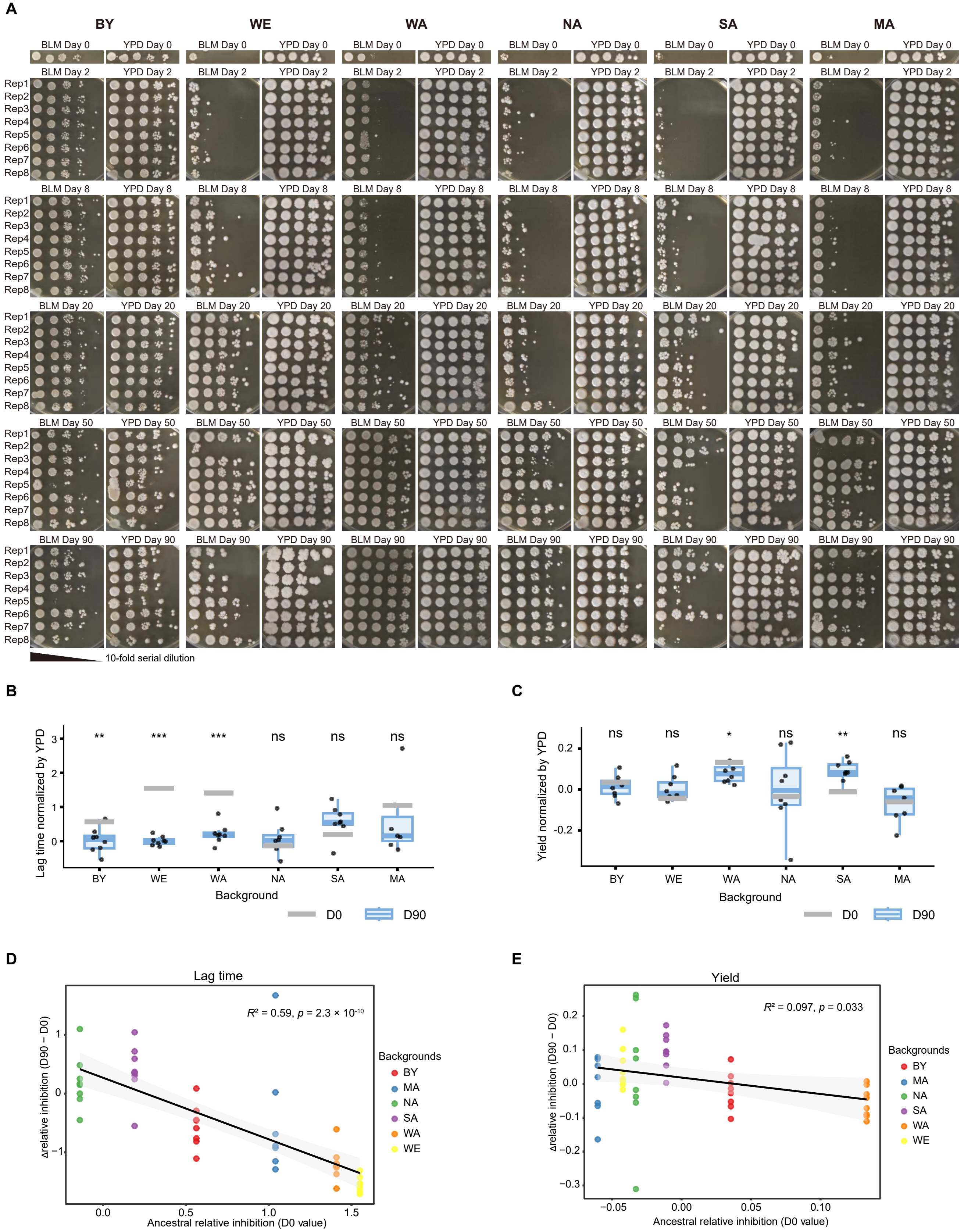
A
BY
WE
WA
NA
SA
MA
BLM Day 0
YPD Day 0
BLM Day 2
YPD Day 2
BLM Day 8
YPD Day 8
BLM Day 20
YPD Day 20
BLM Day 50
YPD Day 50
BLM Day 90
YPD Day 90
Rep1
Rep2
Rep3
Rep4
Rep5
Rep6
Rep7
Rep8
10-fold serial dilution
B
Lag time normalized by YPD
** *** *** ns ns ns
BY WE WA NA SA MA
Background
D0
D90
C
Yield normalized by YPD
ns ns * ns ** ns
BY WE WA NA SA MA
Background
D0
D90
D
Lag time
$R^2$ = 0.59, $p$ = 2.3 × $10^{-10}$
Δrelative inhibition (D90 – D0)
Ancestral relative inhibition (D0 value)
Backgrounds
BY
MA
NA
SA
WA
WE
E
Yield
$R^2$ = 0.097, $p$ = 0.033
Δrelative inhibition (D90 – D0)
Ancestral relative inhibition (D0 value)
Backgrounds
BY
MA
NA
SA
WA
WE

**Figure S1.** Complete phenotypic evolution under bleomycin stress. **(A)** Complete spotting assay results corresponding to the representative images shown in Fig. 1A. Bleomycin-evolved populations were assayed at the indicated time points on YPD plates with or without 12.5 μg/mL bleomycin. Ten-fold serial dilutions were spotted from left to right, with eight biological replicates arranged from top to bottom. **(B-C)** Enhanced bleomycin resistance in evolved populations. We normalized the lag time **(B)** and yield **(C)** in the presence of drug by those in its absence, calculated as (with BLM − without BLM) / without BLM. Statistical significance was determined by comparing each D90 population with the corresponding D0 strain using t-tests. Significance is indicated in the panels: ns, $p > 0.05$; *$p < 0.05$, **$p < 0.01$, and ***$p < 0.001$. **(D-E)** Correlation between initial growth inhibition and growth improvement measured by lag time **(D)** and yield **(E)**. Growth improvement was calculated as the change in growth inhibition between D90 and D0 strains. Each point represents an independently evolved lineage. Pearson correlation coefficients ($R^2$) and corresponding p values are shown in each panel.

**Figure S2**

**A**

| | D0_BY | D90_BY_YPD_1 | D90_BY_YPD_3 | D90_BY_YPD_5 | D90_BY_YPD_7 | D0_WE | D90_WE_YPD_1 | D90_WE_YPD_3 | D90_WE_YPD_5 | D90_WE_YPD_7 | D0_WA | D90_WA_YPD_1 | D90_WA_YPD_3 | D90_WA_YPD_5 | D90_WA_YPD_7 | D0_NA | D90_NA_YPD_1 | D90_NA_YPD_3 | D90_NA_YPD_5 | D90_NA_YPD_7 | D0_SA | D90_SA_YPD_1 | D90_SA_YPD_3 | D90_SA_YPD_6 | D90_SA_YPD_8 | D0_MA | D90_MA_YPD_1 | D90_MA_YPD_3 | D90_MA_YPD_6 | D90_MA_YPD_8 |
|---|---|---|---|---|---|---|---|---|---|---|---|---|---|---|---|---|---|---|---|---|---|---|---|---|---|---|---|---|---|---|
| XVI | 1.9 | 2.0 | 2.0 | 1.9 | 1.9 | 2.0 | 2.0 | 2.0 | 2.0 | 2.0 | 2.0 | 2.0 | 2.0 | 2.0 | 2.0 | 2.0 | 2.0 | 2.1 | 2.1 | 2.0 | 2.0 | 2.0 | 2.0 | 2.0 | 2.0 | 2.0 | 2.0 | 2.0 | 2.0 | 2.0 |
| XV | 1.9 | 2.0 | 2.0 | 1.9 | 2.0 | 2.0 | 2.0 | 2.0 | 2.0 | 2.0 | 2.0 | 1.9 | 1.9 | 2.0 | 1.9 | 2.0 | 2.0 | 2.0 | 2.1 | 2.1 | 2.0 | 2.0 | 2.1 | 2.0 | 2.0 | 2.0 | 2.0 | 2.0 | 2.0 | 2.0 |
| XIV | 2.0 | 2.1 | 2.1 | 2.1 | 2.1 | 2.0 | 2.0 | 2.0 | 2.0 | 2.0 | 2.1 | 2.0 | 2.0 | 2.1 | 2.1 | 2.0 | 2.1 | 2.1 | 1.9 | 2.0 | 2.0 | 2.0 | 2.0 | 2.0 | 2.0 | 2.0 | 2.0 | 2.0 | 2.0 | 2.0 |
| XIII | 2.0 | 2.1 | 2.1 | 2.0 | 2.0 | 2.0 | 2.1 | 2.0 | 2.1 | 2.0 | 2.0 | 2.0 | 2.0 | 2.0 | 2.0 | 2.0 | 2.0 | 2.0 | 2.1 | 2.1 | 2.1 | 2.0 | 2.1 | 2.0 | 2.1 | 2.0 | 2.1 | 2.0 | 2.0 | 2.0 |
| XII | 1.9 | 1.9 | 1.9 | 1.9 | 1.9 | 2.1 | 2.1 | 2.1 | 2.1 | 2.2 | 1.9 | 1.9 | 1.9 | 1.8 | 1.9 | 2.0 | 1.9 | 1.9 | 2.1 | 1.9 | 1.9 | 1.9 | 2.0 | 1.9 | 1.9 | 2.0 | 2.0 | 2.0 | 2.0 | 2.0 |
| XI | 2.1 | 2.1 | 2.1 | 2.1 | 2.1 | 1.9 | 2.0 | 2.0 | 1.9 | 1.9 | 2.0 | 2.1 | 2.1 | 2.1 | 2.1 | 2.0 | 2.1 | 2.0 | 1.9 | 2.0 | 2.0 | 2.0 | 1.9 | 2.0 | 2.0 | 2.0 | 2.0 | 2.0 | 2.0 | 2.0 |
| X | 2.0 | 2.1 | 2.0 | 2.0 | 2.0 | 2.0 | 2.0 | 2.0 | 2.0 | 1.9 | 2.0 | 2.0 | 2.0 | 2.1 | 2.1 | 2.0 | 2.0 | 2.0 | 2.0 | 2.0 | 2.0 | 2.2 | 2.0 | 2.0 | 2.0 | 2.0 | 2.0 | 2.0 | 2.0 | 2.0 |
| IX | 2.2 | 2.2 | 2.2 | 2.2 | 2.2 | 1.9 | 1.9 | 1.9 | 1.9 | 1.9 | 2.2 | 2.1 | 2.2 | 2.2 | 2.2 | 2.0 | 2.3 | 2.0 | 1.7 | 1.9 | 2.1 | 2.2 | 1.9 | 2.3 | 2.5 | 2.1 | 2.1 | 2.1 | 2.1 | 2.1 |
| VIII | 2.1 | 2.1 | 2.1 | 2.1 | 2.0 | 1.9 | 1.9 | 1.9 | 1.9 | 1.9 | 2.2 | 2.1 | 2.1 | 2.2 | 2.2 | 2.0 | 2.0 | 1.9 | 1.8 | 1.9 | 2.0 | 2.1 | 1.9 | 2.2 | 2.1 | 2.0 | 2.0 | 2.0 | 2.0 | 2.0 |
| VII | 1.9 | 2.0 | 2.0 | 2.0 | 2.0 | 2.1 | 2.0 | 2.0 | 2.1 | 2.1 | 1.9 | 1.9 | 1.9 | 1.9 | 1.9 | 2.0 | 2.0 | 2.0 | 2.1 | 2.1 | 2.0 | 2.0 | 2.1 | 2.1 | 2.0 | 2.0 | 2.0 | 2.0 | 2.0 | 2.0 |
| VI | 2.4 | 2.3 | 2.3 | 2.4 | 2.3 | 1.7 | 1.7 | 1.7 | 1.7 | 1.7 | 2.2 | 2.2 | 2.2 | 2.3 | 2.3 | 2.0 | 1.8 | 2.0 | 1.6 | 1.8 | 2.0 | 2.0 | 1.8 | 2.0 | 2.0 | 2.0 | 2.1 | 2.1 | 2.0 | 2.0 |
| V | 2.2 | 2.2 | 2.2 | 2.3 | 2.1 | 2.0 | 2.0 | 2.0 | 2.0 | 2.0 | 2.1 | 2.1 | 2.1 | 2.1 | 2.2 | 2.0 | 1.9 | 2.0 | 1.9 | 2.0 | 2.0 | 2.0 | 1.9 | 2.2 | 2.0 | 2.0 | 2.1 | 2.1 | 2.1 | 2.0 |
| IV | 1.8 | 1.8 | 1.8 | 1.8 | 1.8 | 2.1 | 2.1 | 2.1 | 2.1 | 2.1 | 1.8 | 1.9 | 1.8 | 1.8 | 1.8 | 2.0 | 2.1 | 2.0 | 2.2 | 2.0 | 2.0 | 2.0 | 2.1 | 1.9 | 1.9 | 2.0 | 1.9 | 1.9 | 2.0 | 2.0 |
| III | 2.3 | 1.7 | 1.8 | 2.1 | 2.1 | 1.7 | 1.7 | 1.7 | 1.7 | 1.7 | 2.1 | 2.1 | 2.1 | 2.2 | 2.2 | 1.9 | 1.8 | 1.9 | 1.5 | 1.8 | 2.0 | 2.0 | 1.7 | 1.9 | 1.9 | 1.9 | 2.0 | 2.0 | 1.9 | 2.0 |
| II | 2.1 | 2.1 | 2.1 | 2.1 | 2.1 | 2.0 | 2.0 | 2.0 | 2.0 | 2.0 | 2.0 | 2.0 | 2.0 | 2.1 | 2.1 | 2.0 | 1.9 | 2.0 | 1.9 | 2.0 | 2.0 | 2.0 | 2.0 | 2.0 | 2.0 | 2.0 | 2.0 | 2.0 | 2.0 | 2.0 |
| I | 2.4 | 2.3 | 2.3 | 2.4 | 2.3 | 1.7 | 1.7 | 1.7 | 1.6 | 1.6 | 2.2 | 2.2 | 2.2 | 2.2 | 2.3 | 1.9 | 1.9 | 1.9 | 1.5 | 1.8 | 2.0 | 2.1 | 1.7 | 2.0 | 2.5 | 1.9 | 2.0 | 2.0 | 1.9 | 1.9 |

**B**

| | D0_BY | D90_BY_BLM_6_C1 | D90_BY_BLM_6_C2 | D0_WE | D90_WE_BLM_1_C1 | D90_WE_BLM_1_C2 | D90_WE_BLM_3_C1 | D90_WE_BLM_5_C1 | D90_WE_BLM_5_C3 | D90_WE_BLM_5_C7 | D90_WE_BLM_8_C1 | D90_WE_BLM_8_C2 | D90_WE_BLM_8_C4 | D0_WA | D90_WA_BLM_3_C1 | D90_WA_BLM_3_C2 | D90_WA_BLM_3_C3 | D0_NA | D90_NA_BLM_1_C1 | D90_NA_BLM_1_C2 | D90_NA_BLM_5_C1 | D90_NA_BLM_5_C3 | D90_NA_BLM_7_C2 | D90_NA_BLM_7_C3 | D90_NA_BLM_7_C4 | D90_NA_BLM_8_C2 | D0_MA | D90_MA_BLM_3_C1 | D90_MA_BLM_3_C2 | D90_MA_BLM_5_C1 | D90_MA_BLM_5_C2 | D90_MA_BLM_8_C1 |
|---|---|---|---|---|---|---|---|---|---|---|---|---|---|---|---|---|---|---|---|---|---|---|---|---|---|---|---|---|---|---|---|---|
| XVI | 1.9 | 1.9 | 2.0 | 2.0 | 2.0 | 2.0 | 2.0 | 2.0 | 2.0 | 2.0 | 2.0 | 2.0 | 2.0 | 2.0 | 2.0 | 2.0 | 2.0 | 2.0 | 2.0 | 2.0 | 2.0 | 2.0 | 2.0 | 2.0 | 2.0 | 2.0 | 2.0 | 2.0 | 2.0 | 2.0 | 2.0 | 1.9 |
| XV | 1.9 | 2.0 | 2.0 | 2.0 | 2.1 | 2.0 | 2.0 | 1.9 | 1.9 | 1.9 | 2.0 | 2.0 | 2.0 | 2.0 | 1.9 | 1.9 | 2.0 | 2.0 | 2.0 | 2.1 | 2.0 | 2.1 | 2.0 | 2.0 | 2.0 | 2.0 | 2.0 | 2.0 | 2.0 | 2.0 | 2.0 | 1.9 |
| XIV | 2.0 | 2.0 | 2.1 | 2.0 | 2.0 | 2.0 | 2.0 | 2.1 | 2.1 | 1.9 | 2.0 | 2.0 | 2.0 | 2.1 | 2.1 | 2.0 | 2.1 | 2.0 | 2.0 | 2.0 | 2.0 | 2.0 | 2.0 | 2.0 | 2.0 | 2.0 | 2.0 | 2.1 | 2.1 | 2.0 | 2.1 | 2.0 |
| XIII | 2.0 | 2.0 | 2.1 | 2.0 | 2.1 | 2.0 | 2.0 | 2.0 | 2.0 | 2.0 | 2.0 | 2.0 | 2.0 | 2.0 | 2.0 | 3.0 | 2.0 | 2.0 | 2.0 | 2.0 | 2.0 | 2.0 | 2.0 | 2.0 | 2.0 | 2.0 | 2.0 | 2.1 | 2.1 | 2.1 | 2.1 | 3.0 |
| XII | 1.9 | 1.9 | 1.9 | 2.1 | 2.0 | 1.9 | 2.0 | 1.8 | 1.9 | 1.8 | 1.9 | 1.9 | 2.0 | 1.9 | 1.8 | 1.8 | 1.8 | 2.0 | 1.8 | 2.1 | 2.0 | 2.0 | 1.9 | 1.9 | 1.9 | 1.9 | 2.0 | 1.9 | 1.9 | 1.9 | 2.0 | 1.9 |
| XI | 2.1 | 2.1 | 2.1 | 1.9 | 2.0 | 2.0 | 2.0 | 2.0 | 2.1 | 2.1 | 2.0 | 2.0 | 2.0 | 2.0 | 2.1 | 2.1 | 2.1 | 2.0 | 2.0 | 2.0 | 2.0 | 2.0 | 2.0 | 2.1 | 2.0 | 2.0 | 2.0 | 2.0 | 2.0 | 2.0 | 2.0 | 2.0 |
| X | 2.0 | 2.1 | 2.1 | 2.0 | 2.0 | 2.0 | 2.0 | 2.0 | 2.0 | 2.1 | 2.0 | 2.0 | 2.0 | 2.0 | 2.0 | 2.0 | 2.1 | 2.0 | 2.0 | 2.0 | 2.0 | 2.0 | 2.0 | 2.1 | 2.0 | 2.0 | 2.0 | 2.0 | 2.0 | 2.0 | 2.0 | 2.0 |
| IX | 2.2 | 2.2 | 2.2 | 1.9 | 2.0 | 2.2 | 2.1 | 2.1 | 2.2 | 2.2 | 2.1 | 2.1 | 2.0 | 2.2 | 2.2 | 2.2 | 2.1 | 2.0 | 2.1 | 1.9 | 2.0 | 2.0 | 2.0 | 2.1 | 2.1 | 2.0 | 2.1 | 2.2 | 2.2 | 2.2 | 2.2 | 3.1 |
| VIII | 2.1 | 2.0 | 2.1 | 1.9 | 1.9 | 2.0 | 2.0 | 3.1 | 2.1 | 2.0 | 2.0 | 2.0 | 2.0 | 2.2 | 2.1 | 2.1 | 2.1 | 2.0 | 2.0 | 1.8 | 2.0 | 2.0 | 2.0 | 2.0 | 2.0 | 2.0 | 2.0 | 2.1 | 2.1 | 2.1 | 2.1 | 2.0 |
| VII | 1.9 | 2.0 | 2.0 | 2.1 | 2.0 | 2.0 | 2.0 | 2.0 | 2.0 | 2.0 | 2.0 | 2.0 | 2.0 | 1.9 | 1.9 | 1.9 | 2.0 | 2.0 | 1.9 | 2.1 | 2.0 | 2.0 | 2.0 | 2.0 | 2.0 | 2.0 | 2.0 | 2.1 | 2.0 | 2.1 | 2.0 | 2.0 |
| VI | 2.4 | 2.3 | 2.3 | 1.7 | 1.8 | 2.1 | 2.0 | 2.1 | 2.1 | 2.2 | 2.0 | 2.0 | 2.0 | 2.2 | 2.2 | 2.2 | 2.1 | 2.0 | 2.1 | 1.8 | 1.9 | 2.0 | 2.0 | 2.1 | 2.0 | 2.0 | 2.0 | 2.2 | 2.1 | 2.2 | 2.1 | 2.0 |
| V | 2.2 | 2.1 | 2.2 | 2.0 | 2.0 | 2.1 | 2.1 | 2.1 | 2.1 | 2.2 | 2.1 | 2.1 | 2.0 | 2.1 | 2.1 | 2.1 | 2.1 | 2.0 | 3.0 | 1.9 | 2.0 | 2.0 | 2.0 | 2.1 | 2.1 | 2.0 | 2.0 | 2.2 | 2.1 | 2.2 | 2.1 | 3.0 |
| IV | 1.8 | 1.9 | 1.8 | 2.1 | 2.1 | 1.9 | 2.0 | 1.9 | 1.9 | 1.9 | 2.0 | 2.0 | 2.0 | 1.8 | 1.8 | 1.8 | 1.9 | 2.0 | 1.9 | 2.1 | 2.0 | 2.0 | 2.0 | 1.9 | 2.0 | 2.0 | 2.0 | 1.8 | 1.9 | 1.8 | 1.9 | 1.9 |
| III | 2.3 | 2.2 | 2.2 | 1.7 | 1.8 | 2.0 | 1.9 | 2.0 | 2.1 | 2.2 | 2.0 | 2.0 | 2.0 | 2.1 | 2.2 | 2.1 | 2.1 | 1.9 | 2.1 | 1.8 | 1.8 | 1.9 | 2.0 | 2.0 | 2.0 | 2.0 | 1.9 | 2.1 | 2.1 | 2.2 | 2.0 | 2.0 |
| II | 2.1 | 2.0 | 2.1 | 2.0 | 2.0 | 2.1 | 2.0 | 2.1 | 2.0 | 2.0 | 2.0 | 2.0 | 2.0 | 2.0 | 3.0 | 2.0 | 2.0 | 2.0 | 2.9 | 2.0 | 2.0 | 2.0 | 2.0 | 2.0 | 2.0 | 2.0 | 2.0 | 2.0 | 2.0 | 2.0 | 2.0 | 2.0 |
| I | 2.4 | 2.3 | 2.1 | 1.7 | 1.7 | 2.0 | 1.9 | 2.0 | 2.1 | 2.2 | 2.0 | 2.0 | 2.0 | 2.2 | 2.2 | 2.2 | 2.1 | 1.9 | 3.1 | 1.8 | 1.9 | 1.9 | 2.0 | 2.0 | 2.0 | 2.0 | 1.9 | 2.2 | 2.1 | 2.2 | 2.0 | 1.9 |

**Figure S2.** Aneuploidy profiles of YPD-evolved populations and the clones isolated from the bleomycin-evolved populations. **(A)** YPD-evolved populations. **(B)** The clones isolated from the bleomycin-evolved populations. Chromosome copy number = 2 × (median depth of the chromosome / median depth of the genome).

Figure S3

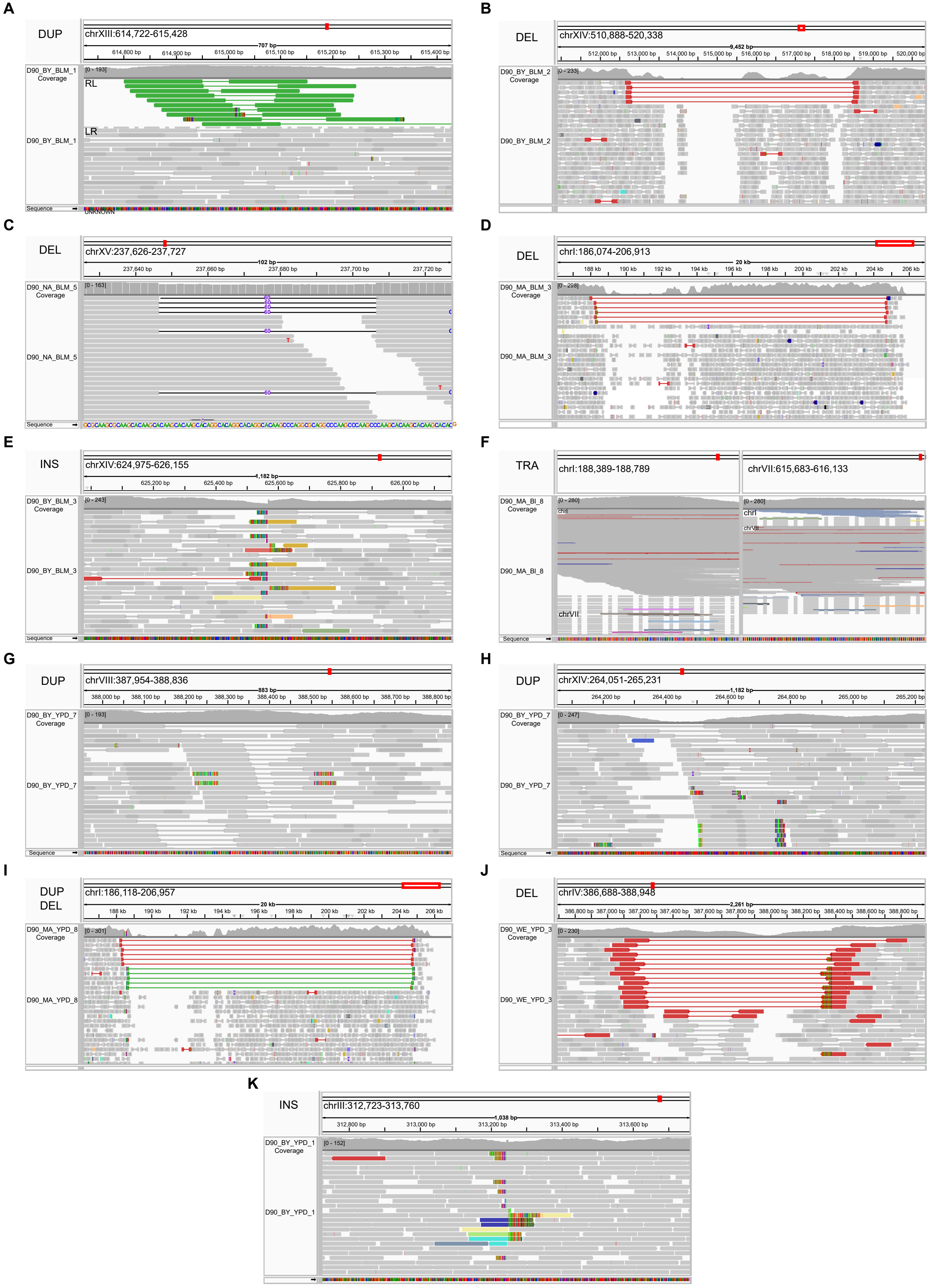
A
DUP
chrXIII:614,722-615,428
D90_BY_BLM_1 Coverage
RL
LR
D90_BY_BLM_1
B
DEL
chrXIV:510,888-520,338
D90_BY_BLM_2 Coverage
D90_BY_BLM_2
C
DEL
chrXV:237,626-237,727
D90_NA_BLM_5 Coverage
D90_NA_BLM_5
D
DEL
chrI:186,074-206,913
D90_MA_BLM_3 Coverage
D90_MA_BLM_3
E
INS
chrXIV:624,975-626,155
D90_BY_BLM_3 Coverage
D90_BY_BLM_3
F
TRA
chrI:188,389-188,789
chrVII:615,683-616,133
D90_MA_BI_8 Coverage
D90_MA_BI_8
chrI
chrVII
G
DUP
chrVIII:387,954-388,836
D90_BY_YPD_7 Coverage
D90_BY_YPD_7
H
DUP
chrXIV:264,051-265,231
D90_BY_YPD_7 Coverage
D90_BY_YPD_7
I
DUP
DEL
chrI:186,118-206,957
D90_MA_YPD_8 Coverage
D90_MA_YPD_8
J
DEL
chrIV:386,688-388,948
D90_WE_YPD_3 Coverage
D90_WE_YPD_3
K
INS
chrIII:312,723-313,760
D90_BY_YPD_1 Coverage
D90_BY_YPD_1

**Figure S3.** IGV visualization of structural variants identified in evolved populations. **(A-K)** IGV screenshots showing read alignments supporting 12 structural variants identified in BLM- and YPD-evolved populations. Split reads and discordant read pairs supporting the breakpoints are shown. Sample names, structural variant types, and genomic coordinates are indicated in each panel.

**Figure S4**

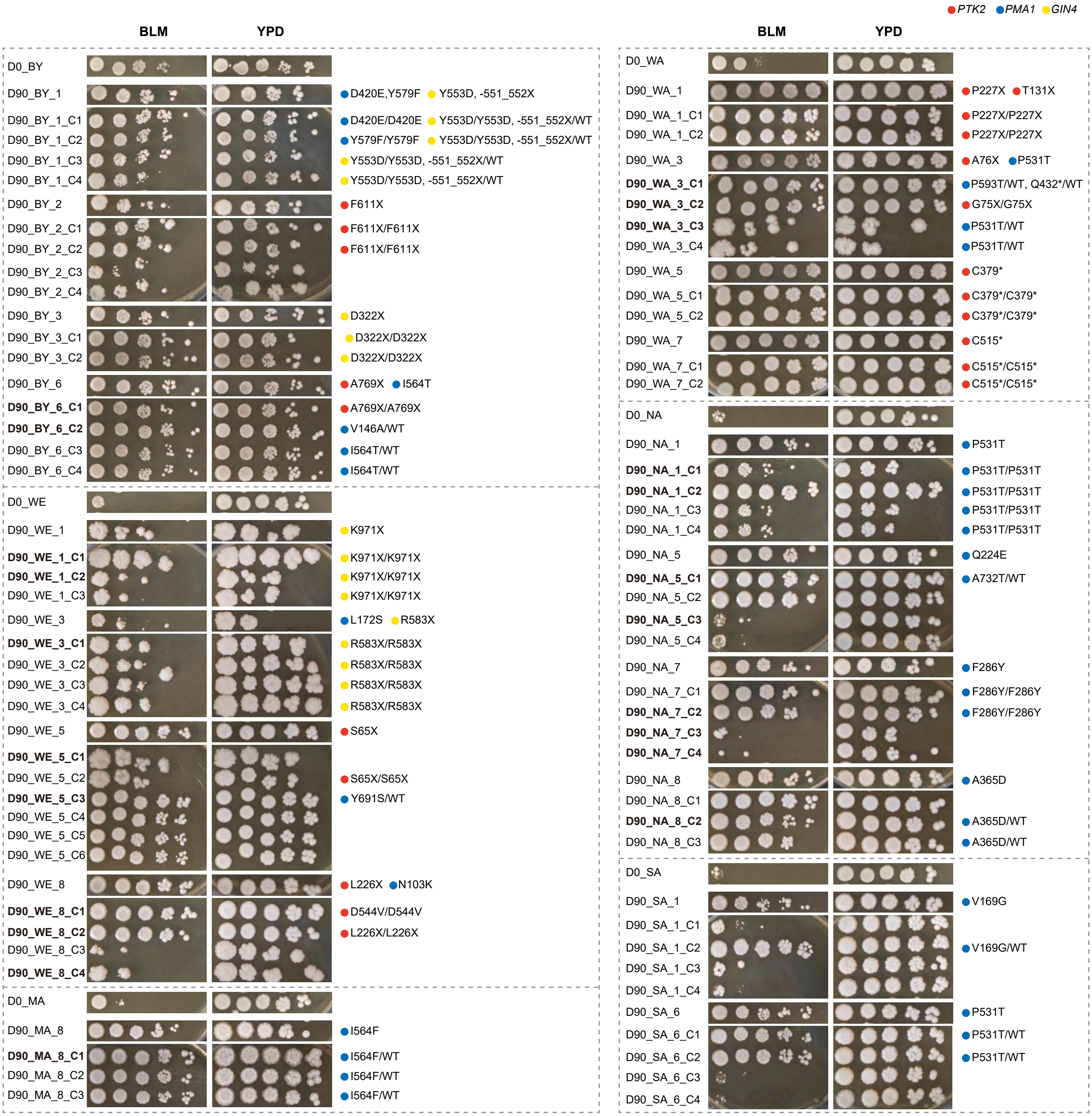


**Figure S4.** Phenotypic validation of populations and clones with *PMA1*, *PTK2*, and *GIN4* mutations. Samples shown in bold were subjected to whole-genome sequencing at the clonal level.

**Figure S5**

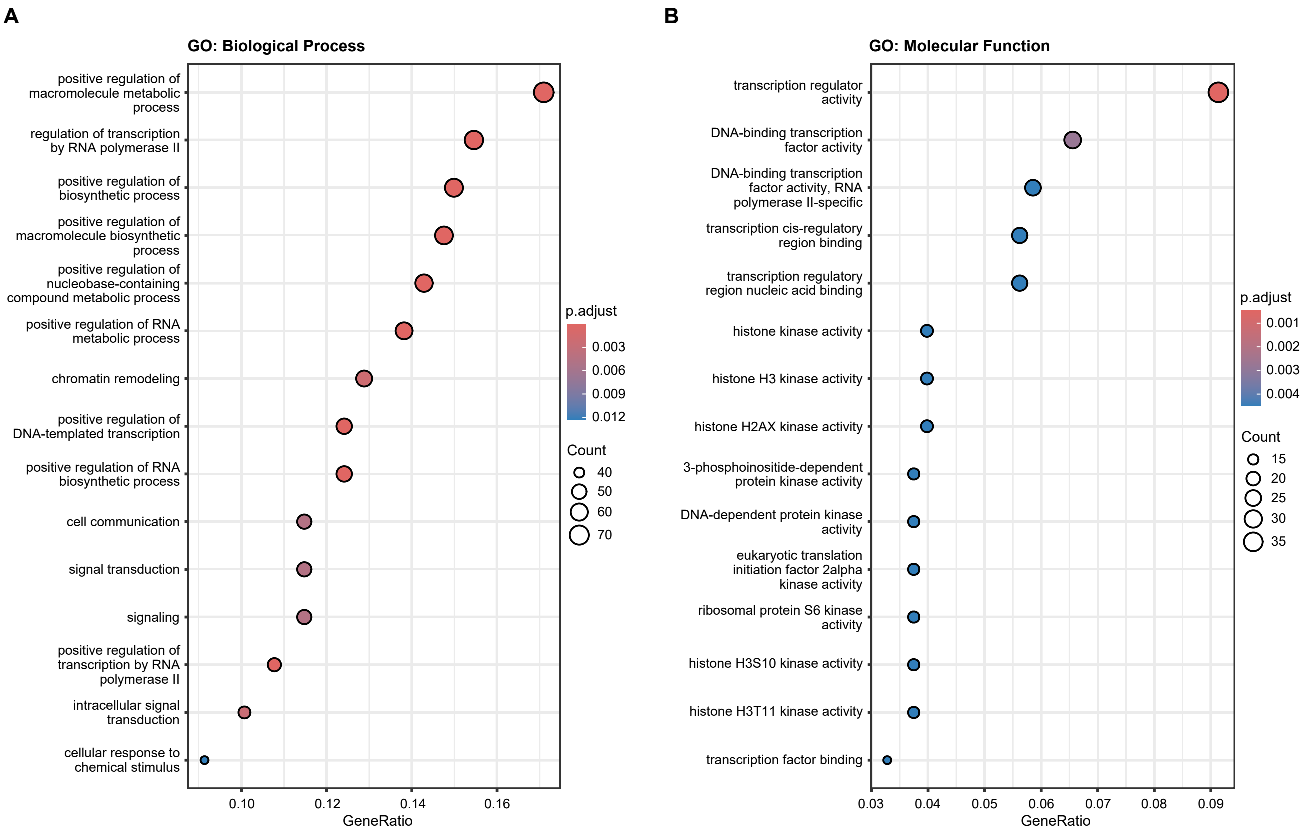


**Figure S5.** GO analysis of bleomycin-specific mutations with moderate or high impact. **(A)** Biological process terms. **(B)** Molecular function terms.

**Figure S6**

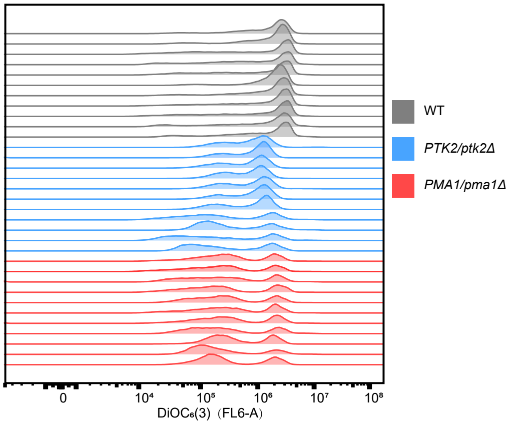


**Figure S6.** Plasma membrane potential measurement by DiOC6(3) staining and flow cytometry.
